# Disrupted Forward Connectivity in Parieto-temporal Network Impairs Memory Performance in Alzheimer’s Disease

**DOI:** 10.1101/2025.06.05.658022

**Authors:** Yanin Suksangkharn, Björn Hendrik Schott, Peter Zeidman, Niklas Vockert, René Lattmann, Hartmut Schütze, Renat Yakupov, Oliver Peters, Julian Hellmann-Regen, Lukas Preis, Ersin Ersözlü, Josef Priller, Eike Jakob Spruth, Maria Gemenetzi, Anja Schneider, Klaus Fliessbach, Jens Wiltfang, Claudia Bartels, Ayda Rostamzadeh, Wenzel Glanz, Enise I. Incesoy, Stefan Teipel, Ingo Kilimann, Doreen Goerss, Christoph Laske, Annika Spottke, Marie Kronmüller, Frederic Brosseron, Falk Lüsebrink, Matthias Schmid, Luca Kleineidam, Melina Stark, Stefan Hetzer, Peter Dechent, Klaus Scheffler, Frank Jessen, Anne Maass, Emrah Düzel, Gabriel Ziegler

## Abstract

Alzheimer’s disease (AD) is characterised by the accumulation of beta-amyloid (Aβ) and tau proteins, leading to neurodegeneration and cognitive decline. While Aβ and tau are known to disrupt synaptic function, the mechanisms linking these molecular pathologies to network-level dysfunction and memory impairment remain poorly understood. Here we investigated the effects of Aβ and tau pathology (CSF Aβ42/40 ratio and tau phosphorylated at position 181, p-tau-181, respectively) on effective connectivity (EC) related to memory encoding, which may constitute a link between synaptic pathology and cognitive outcomes. Functional magnetic resonance imaging (fMRI) during visual memory encoding was acquired from participants of the multicentric DZNE Longitudinal Cognitive Impairment and Dementia Study (DELCODE), including 203 cognitively normal older participants (CN) as well as individuals with subjective cognitive decline (SCD; N = 204), mild cognitive impairment (MCI; N = 65), and early dementia due to AD (DAT; N = 21). EC was assessed by applying Dynamic causal modelling (DCM) to the fMRI data, using brain regions previously implicated in memory-encoding: the parahippocampal place area (PPA), the hippocampus (HC) and the precuneus (PCU). Disruptions in forward connectivity from the PPA to the HC and PCU were associated with both memory impairment and indices of AD pathology. Specifically, reduced excitatory EC from the PPA to the HC was associated with higher p-tau-181 levels and correlated with poorer memory performance. Diminished inhibitory EC from the PPA to the PCU was driven by both tau and amyloid pathology and was likewise linked to memory decline. Our findings suggest that disrupted forward connectivity within the temporo-parietal memory network constitutes a candidate mechanism mediating the relationship between molecular pathology and cognitive dysfunction.

## Introduction

As Alzheimer’s disease (AD) anatomically spreads in the brain, it causes cognitive impairment and decline due to neurodegeneration and synaptic dysfunction. Impairment and decline are not only related to which brain regions are affected but also to how different brain regions interact with each other. Here we investigate how excitatory and inhibitory interactions between brain regions of the episodic memory network relate to impairment and decline in AD.

AD is characterised by the accumulation of beta-amyloid (Aβ) plaques and tau protein aggregates, both contributing synergistically to synaptic dysfunction, neuronal loss, and cognitive decline.^1,2^ Aβ accumulation leads to extracellular plaque formation, impairing neuronal communication through disruptions of neurotransmission, neuroinflammation, and mitochondrial dysfunction.^3–6^ Moreover, Aβ accumulation exacerbates tau hyperphosphorylation, further driving neurodegeneration.^3,7^ Hyperphosphorylated tau independently forms intracellular aggregates, progressing into neurofibrillary tangles that impair axonal transport, synaptic integrity, and neuronal viability, ultimately manifesting as cognitive impairment.^8,9^

Aβ accumulation begins in neocortical areas, particularly within associative networks, before progressing to deeper brain structures.^10,11^ Tau pathology follows a different anatomical spreading pattern: it often originates in the brainstem and spreads to limbic regions critical for memory and spatial navigation, most notably the entorhinal cortex and hippocampal formation.^12,13^ Recent evidence suggests that Aβ-induced hyperconnectivity facilitates the spread of tau pathology across functionally connected brain regions, especially to Aβ-accumulating regions, further exacerbating synaptic disruption and cognitive decline.^14–16^ The distinct spatial trajectories and directionally asymmetric interactions contribute to region-specific vulnerabilities.^17^ Understanding how these pathologies interact with network-level dynamics of human memory function can provide insights into the neural mechanisms linking AD pathology to clinical manifestation.

Functional magnetic resonance imaging (fMRI), particularly in combination with measures of brain connectivity, provides a powerful tool to investigate how AD pathology may impact neural network dynamics during memory processes. Effective connectivity (EC) stands out among other functional connectivity measures by reflecting the causal influence one brain region exerts on another, thereby providing directional information.**^18,19^**Dynamic causal modelling (DCM) is a widely used approach to assess EC of fMRI data and integrates biologically plausible properties, such as non-linear interactions and biophysical constraints, into the modelling of EC.**^20,21^**At the scale of fMRI, EC can be interpreted as an indicator of network-level neuronal interactions, rendering it a promising tool for investigating the link between molecular pathology and memory impairment across the pre-clinical stages of AD.

In the present study, we employed DCM to investigate how network dysfunction mediates the link between Alzheimer’s disease pathology and cognitive decline. Based on the commonly employed amyloid-tau-neurodegeneration (ATN) framework^1,22,23^, we focused on the effects on ATN proxies, namely amyloid pathology (CSF Aβ42/40 ratio), tau (CSF p-tau-181), and hippocampal volume on EC within the memory-encoding network, which is prominently affected by AD pathology.^24^ Next, we aimed to assess a potential association of altered EC with episodic memory deficits in pre-clinical Alzheimer’s disease. We analysed fMRI acquired in a cohort of 493 older adults across the AD risk spectrum, including cognitively normal older adults (CN), individuals with subjective cognitive decline (SCD) and with mild cognitive impairment (MCI) as well as patients with early-stage dementia of Alzheimer’s type (DAT). DCM was applied to fMRI data acquired during a visual memory-encoding task, which has previously been evaluated in individuals at risk for AD.^25–27^ Based on a previous DCM study using the same paradigm in an independent cohort^28^, we chose the hippocampus (HC), the parahippocampal place area (PPA), and the precuneus (PCU) as regions of interest. We hypothesised that EC between these would be modulated by measures of both amyloid and tau burden pathology as well as their interactions. Additionally, we determined to explore relationship between hippocampal volume loss as a proxy for neurodegeneration and connectivity alterations.

## Materials and methods

The rationale, procedures and methods for our study are summarized in Figure 1.

**Figure 1.**
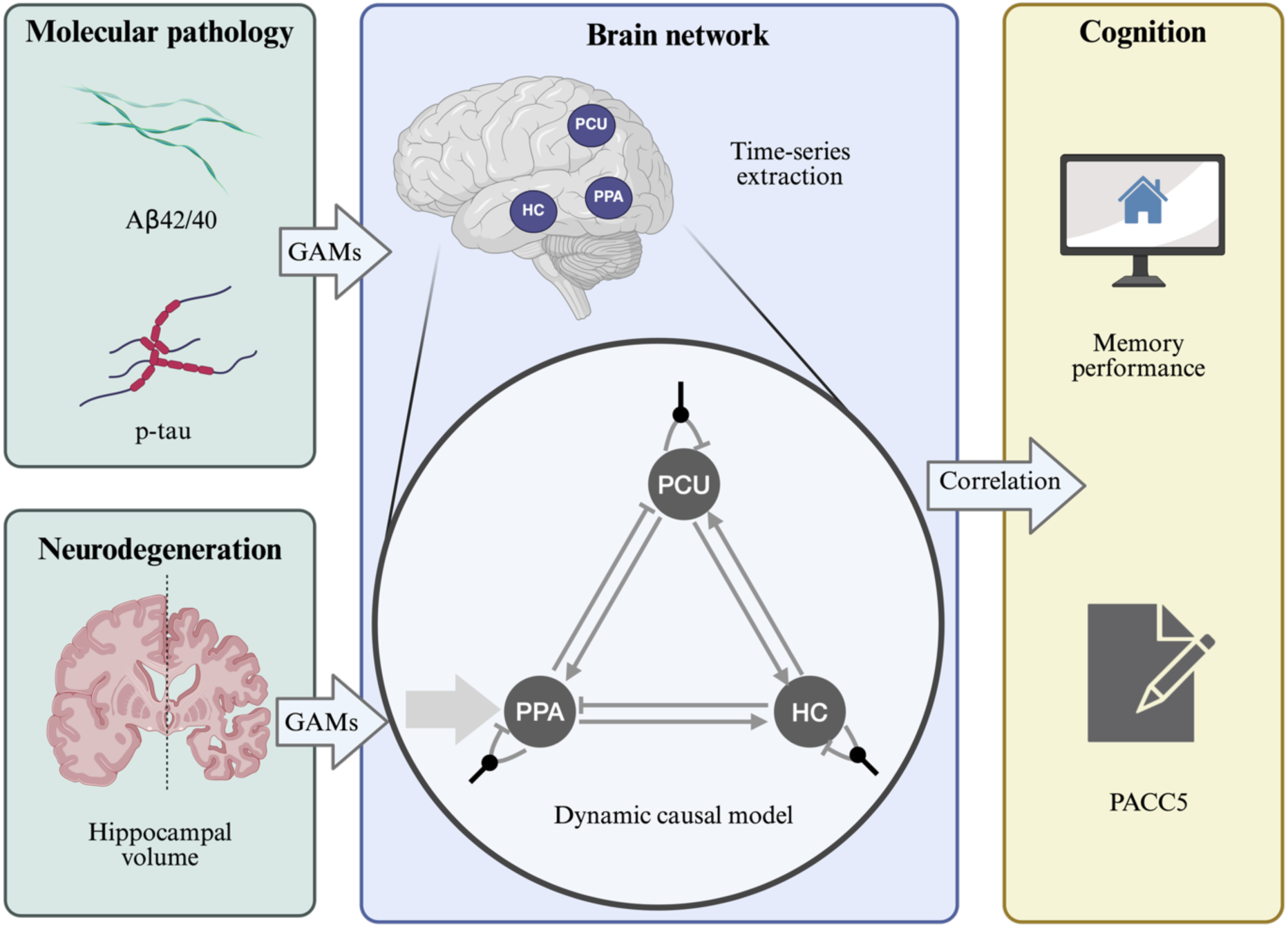
Schematic overview of the study procedures. (Left) Biomarkers of Alzheimer’s disease include (1) molecular pathology, such as the Amyloid-β42/40 ratio (Aβ42/40) and phospho-tau181 (p-tau-181) from cerebrospinal fluid (CSF), and (2) neurodegeneration, represented by hippocampal volume. Generalised additive models (GAMs) were used to investigate the relationship between these biomarkers and brain network dynamics. (Middle) Brain network dynamics during memory encoding are represented by a DCM derived from brain activity time-series. Regions of interest include the parahippocampal place area (PPA), hippocampus (HC), and precuneus (PCU). Thin arrows represent intrinsic connectivity between regions, blunt-ended arrows indicate self-inhibitory intrinsic connectivity, and circle-headed arrows denote modulatory influences associated with subsequent memory scores, reflecting successful memory encoding. The thick arrow illustrates the driving input of novel images to the PPA. (Right) Cognition, as the clinical presentation, is represented by memory performance during the task-based fMRI session and the PACC5 score. The correlation between connectivity parameters and cognition was evaluated.

### Study cohort

The study sample comprised of participants of the DZNE Longitudinal Cognitive Impairment and Dementia Study (DELCODE) study, a memory clinic-based multi-center cohort conducted by the German Centre for Neurodegenerative Diseases (DZNE) (for details, see Jessen et al.^29^). Among the 1,079 participants enrolled, 558 individuals had completed the fMRI visual memory-encoding task. Following outlier removal and quality control procedures (see Participant Exclusion), the final sample comprised 493 participants, of whom 235 had available CSF biomarker data.

The study protocol was approved by the local institutional review boards of all participating sites. All participants provided written informed consent prior to inclusion in the study in accordance with the Declaration of Helsinki. The DELCODE study was retrospectively registered with the German Clinical Trials Register (DRKS00007966) on 04 May 2015. Details on data handling and quality control have been previously described.^29^

### Experimental paradigm

Participants underwent fMRI scanning while performed a visual novelty and encoding task, viewing 44 novel indoor scenes, 44 novel outdoor scenes, and 44 repetitions of pre-familiarised images. Participants classified each image as ‘indoor’ or ‘outdoor’ scene via button press. Each stimulus was displayed for 2500 ms, followed by fixation with an optimised jitter ranging from 0.70 s to 2.65 s.^30^ The task lasted approximately 11 minutes, during which 206 functional volumes were recorded. After a 90-minute delay, a computer-based recognition memory test was conducted outside the scanner to assess recognition for the novel images seen in the fMRI session. The test consisted of 88 exposed novel images and two pre-familiarised images, plus 44 new images, using a 5-step response scale where 1 is definitely new to 5 is definitely old. The recognition responses were converted into parametric modulator with an arcsine-transformation

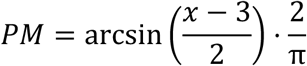

as this approach has previously been demonstrated to outperform categorical designs.^31^ The responses were used as predictor during first-level analysis (denoted as subsequent memory score).

Memory performance of each participant was quantified based on the area under the curve (AUC) of hits (i.e., correctly recognised novelty stimuli) against false alarms (i.e., incorrectly recognised novelty stimuli) by taking the subsequent memory score into account (hence termed A’; for details, see Soch et al.^32^). An A’ of 0.5 reflects pure guessing while Memory performance of 1 reflects perfect performance.

### Characterization of Alzheimer’s pathology and memory performance

Alzheimer’s disease pathology was characterized according to the NIA-AA research framework^23^, encompassing amyloid accumulation (A), pathological tau accumulation (T), and neurodegeneration (N). Amyloid and tau pathology were assessed using cerebrospinal fluid (CSF) biomarkers: the amyloid-β42/40 ratio (Aβ42/40) and phosphorylated tau (p-tau-181), respectively. Neurodegeneration was approximated by adjusted hippocampal volumes derived from MRI-based hippocampal volume adjusted for age, sex, education, total intracranial volume (TIV), and total white matter hyperintensity (WMH) volume. ATN classification was applied using cut-offs determined from the DELCODE dataset via Gaussian mixture modelling: Aβ42/40 ≤ 0.08 pg/ml for A+, p-tau ≥ 73.65 pg/ml for T+, and ≤ 2821 *μl* for N+ (see Düzel et al.^33^ and Heinzinger et al.^34^).

Cognitive outcome was assessed using memory performance (A’) from the memory-encoding fMRI task (see Experimental Paradigm). Additionally, the Preclinical Alzheimer’s Cognitive Composite 5 (PACC5)^35^ was included as an independent validation measure. PACC5 is a composite score, composed of the MMSE, the summed Free and Cued Selective Reminding Test (FCSRT) free and total recall, the Wechsler Memory Scale – Fourth Edition (WMS-IV) Logical Memory Story B delayed recall, the Symbol-Digit Modalities Test (SDMT), and the sum of two category fluency tasks, which are known to predict early disease progression independently. This feature makes PACC5 particularly sensitive to detecting preclinical changes within the AD spectrum.^36^

### MRI acquisition and fMRI data preprocessing

MRI data were acquired using 3T Siemens scanners at the participating sites, including TIM Trio, Verio, Skyra, and Prisma systems. Structural images included T1-weighted (1 *mm*^3^ isotropic resolution) and T2-weighted, optimised for medial temporal lobe volumetry. Functional images were obtained using a T2*-weighted echo-planar imaging (EPI) sequence (TR = 2580 ms, TE = 30 ms, voxel size = 3.5 mm isotropic).

Preprocessing of fMRI data was conducted using SPM12 and involved multiple steps: correction for acquisition delay (slice-timing), head motion (realignment), and magnetic field inhomogeneities (unwarping; using individual phase and magnitude fieldmaps); followed by normalisation into a standard stereotactic reference frame (Montreal Neurological Institute, MNI) and spatial smoothing (Gaussian kernel, FWHM = 6 mm).

### Participant Exclusion

To ensure the reliability of the experiment and account for potential compromises in participant cooperativeness, we excluded extreme outliers based on the following criteria: (1) more than 8 errors in indoor/outdoor judgments during the fMRI; (2) an absolute response bias greater than 1.5 in the post-fMRI recognition memory test; and (3) framewise displacement (FD) exceeding 0.5 mm for any single frame or 0.2 mm for at least 2% of the total frames during the fMRI session. This led to the exclusion of 65 participants (17 CN, 18 SCD, 18 MCI, and 12 DAT), representing 11.6% of the originally tested participants.

### Dynamic Causal Modelling (DCM)

To investigate ATN pathology-related connectivity changes in the temporo-parietal network in the context of a visual memory-encoding paradigm, we implemented Dynamic Causal Modelling (DCM; Figure 1 middle) using SPM12, following the framework described by Zeidman et al.^21^. DCM was employed to estimate EC parameters at the individual level using a Bayesian approach.

**Volume-of-interest (VOI) time-series extraction:** Time-series extraction was performed for the right PPA, HC, and PCU. The boundaries of each VOI were defined as the overlapping area between anatomical and functional constraints. Anatomical constraints were based on masks from the Automated Anatomical Labelling (AAL) atlas.**^37^** Functional constraints were defined using positive memory contrasts for PPA and HC**^32,38–40^**, while the negative memory contrast was used for PCU due to its deactivation pattern during novelty processing and successful memory encoding.**^25,41,42^**Functional constraints were derived from group-level memory contrasts thresholded at *p* < 0.05 (FWE corrected, controlled for amyloid status, p-tau levels, adjusted hippocampal volume, age, sex, and education). Individual-level voxel thresholding was not applied due to subgroup-dependent variability of available voxels in relation to hippocampal volume (*p* < 0.05) and memory performance (*p* < 0.05). Time-series data were adjusted to regress out known confounds and retain the effects of interest (EOI), namely the regressors representing the novel images, their parametric modulation with the subsequent memory scores**^27,32^**, and the pre-exposed images. Time-series from each brain region were summarised in terms of their first principal component across voxels, which formed the data entering the DCM analysis.

**Model specification:** The model was designed to align with the visual memory-encoding paradigm (Figure 1 middle). The brain regions of interest (ROI) included the PPA, the HC, and the PCU. These ROIs were chosen based on earlier studies demonstrating their pronounced role in successful encoding^43^ and their sensitivity to changes in relation to aging^41^ and AD pathology^25,27^. Moreover, DCM using ROIs has previously been successfully applied to the same visual memory-encoding paradigm in independent cohorts of healthy older adults.^28^ Functional time-series of these regions were assumed to be bidirectionally inter-connected and subject to inhibitory self-connectivity. The resulting model space further allowed for each region’s self-inhibition to be modulated by the arcsine-transformed recognition memory responses of each stimulus, serving as a proxy for successful memory encoding on a trial-by-trial basis. The presentation of novel stimuli was included as the driving input to the PPA in our model space.

**Model estimation:** Model estimation was performed using DCM estimation function as implemented in SPM12.**^21^** At the single-subject level, the DCM model was estimated, which included HC, PPA, and PCU, allowing for full connectivity among brain regions. Novelty (i.e., presentation of novel, but not pre-familiarized images) was chosen as driving input to the PPA, and encoding success (the arcsine-transformed recognition memory responses representing encoding success) was included as a potential contextual modulator at all auto-inhibitory self-connections (Figure 2C). Neural responses to the pre-familiarized images formed the implicit baseline.**^28^** Model estimation was performed using variational Laplace**^44,45^** which provides both posterior connectivity estimates and the free energy approximation to the marginal likelihood, which is a score for the quality of the model.

**Figure 2.**
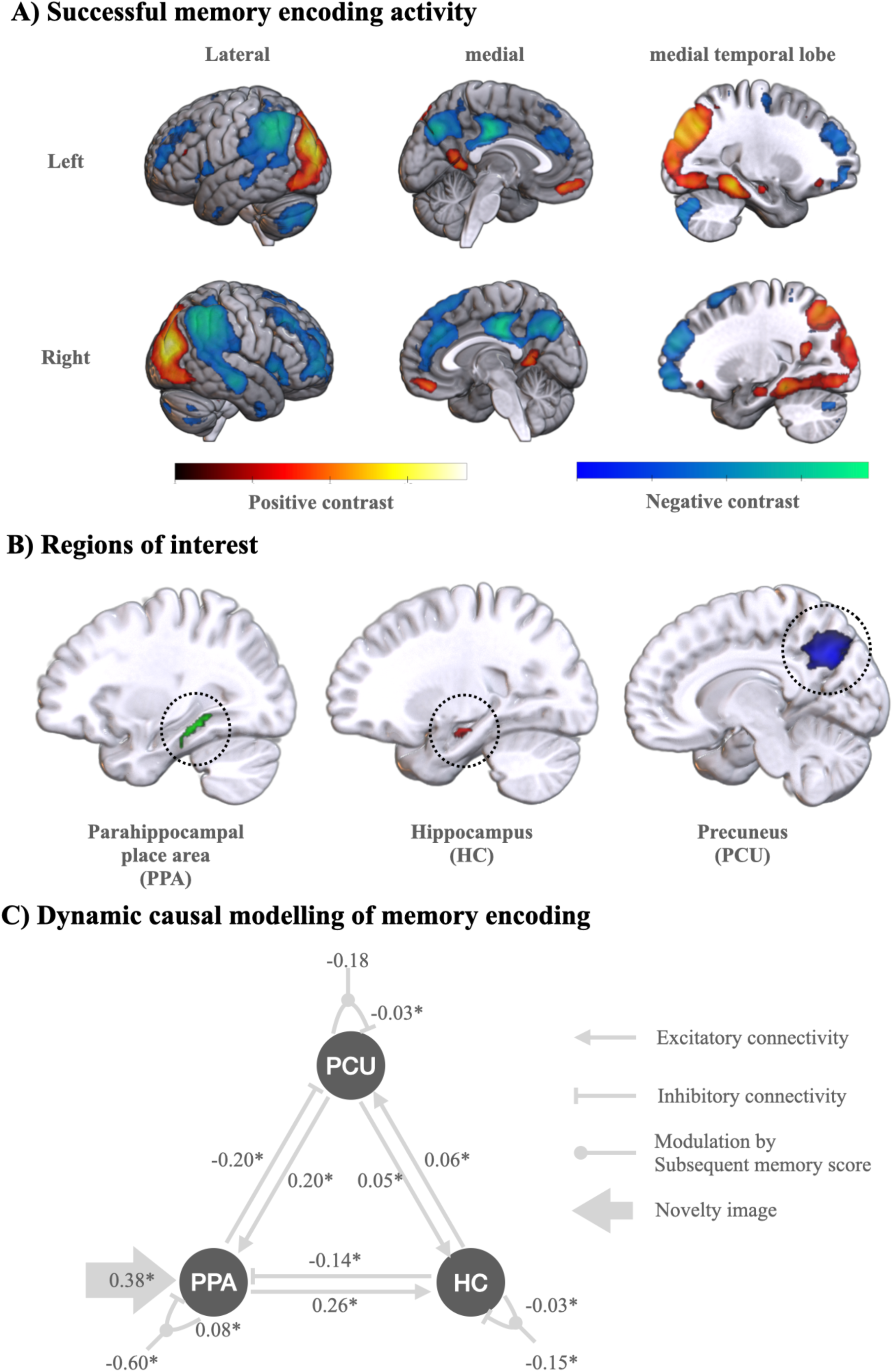
Dynamic Causal Modelling of Memory Encoding. (A) Successful memory-encoding contrast indicating fMRI BOLD signal activations (red/yellow) and deactivations (blue/green) during the encoding of a visual scene that was later successfully remembered. The contrast was derived from parametric regressor for subsequent memory score and was thresholded at *p* < 0.05, corrected for family-wise error (FWE), and controlled for Alzheimer’s disease pathology and covariates. (B) DCM Regions of interest used for time-series data extraction obtained from overlapping significant successful memory-encoding contrast (p<0.05 FWE) from (A) and anatomically defined mask (see methods). **(C)** Dynamic causal modelling of memory-encoding task fMRI in 203 cognitively normal older participants (ages 60 – 80 years), displaying obtained effective connectivity coefficients. Asterisks (*) indicate significant deviations from zero (Wilcoxon rank-sum test with False Discovery Rate (FDR) correction). Abbreviations: PPA (parahippocampal place area), HC (hippocampus), PCU (precuneus).

### Identifying associations of connectivity and memory performance

We first conducted an exploratory analysis of individual DCM parameters with memory performance in the fMRI task itself (Figure 1 right). Due to the observed non-linearity in the data (Supplementary Figure 1), Spearman’s rank correlation was employed. Next, we computed correlations with the PACC5 score was used as an independent measure of memory performance.

To identify DCM parameters with a meaningful relationship to memory performance, we included those with *p* < 0.05 and |*ρ*| > 0.2. P-values were corrected for multiple comparisons using the false discovery rate (FDR) correction.^46^ To verify whether the relationship between EC and memory performance would also hold for cognitive tests independent of the fMRI experiment, we performed out-of-sample estimations of memory performance and PACC5 using intrinsic and modulatory connectivity parameters derived from DCM. Furthermore, leave-one-out cross validation of the model was performed as described for the parametric empirical Bayes (PEB) framework in SPM12.^47^ Afterwards, we evaluated the predictability of the obtained connectivity parameters for memory performance and PACC5 score by using Spearman’s rank correlations controlled for FDR.

### Evaluating effects of AD pathology on effective connectivity

We next investigated potential effects of Aβ42/40 and p-tau on the connectivity estimates that showed significant correlations with memory performance (see above; Figure 1 left). To study potential non-linear relationships (as observed in a previous study**^25^**), a flexible generalised additive model (GAM, pygam**^48^**) was employed. The model can be described as follows:

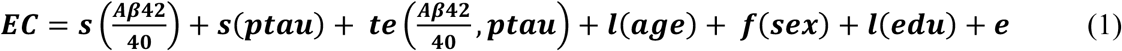

where *s(x)* represents an unknown smooth function of *x*, *te(x)* represents a tensor product term*, l(x)* represents a linear term, and *f(x)* represents a factor (categorical) term. To further investigate a related hypothesis whether functional network alterations could be explained by structural atrophy, additional models were evaluated including *s(h_vol)* as a function of hippocampal volume and covariates as follows:

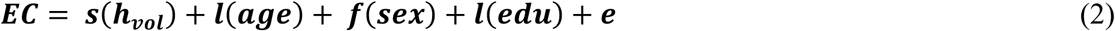

The effect of each term was evaluated using partial dependence plots, which illustrate the isolated (bivariate) effect of each variable on the connectivity estimate. P-values derived from the relationships between biomarkers and EC were adjusted for multiple comparisons using FDR. Note that diagnosis and its interaction with biomarkers were not included in the model to avoid introducing additional.

### Synthesising brain activity with AD pathology

We investigate altered functional brain dynamics during memory encoding in the presence of amyloid and tau pathology by simulation brain activity of 235 individuals in different pathological settings. We utilised the generative properties of the DCM framework, which allow it to simulate and predict network activity and activity signal (BOLD) based on given individual connectivity profile. Using the relationships from Eq. 1, synthetic connectivity profiles were generated from GAMs for four AD pathology groups: A-T-, A+T-, A-T+, and A+T+. Positive and negative status for amyloid and tau were uniformly represented by the mean biomarker levels from CN and DAT groups respectively, i.e., the mean Aβ42/40 ratio from CN for A- and from DAT for A+, and similarly for T status. Furthermore, age and education of all individuals were assigned at the overall sample mean, while sex remained original for all subjects (as in supplementary table 1). The mean biomarker levels of CN and DAT were also corresponding with the ATN classification criteria used in previous studies.^29,33,34^ This approach ensured that biomarker levels remained within a physiologically relevant range, avoiding extreme values that could compromise the accuracy of effect estimation.

We synthesised ROI-based BOLD time courses based on the presence of novel activity and individual responses observed during task-based fMRI sessions, using connectivity profiles from four AT categories. To achieve this, we applied the simulation function from SPM12 (*spm_dcm_simulate.m*) to a synthetic connectivity profile and actual responses from the memory-encoding experiment, generating BOLD signals the three regions (PPA, HC, and PCU) with a signal-to-noise ratio of 1. ROI activity time courses were evaluated by contrasting the synthesized BOLD signals with parametric subsequent memory regressor associated with novelty regressor, incorporating the canonical hemodynamic response function (HRF). Positive contrast was applied to PPA and HC, while negative contrast was applied to PCU, reflecting expected physiological response patterns seen in group level activation contrasts.^25^ Comparisons among the three ROIs were performed using the Wilcoxon rank test with FDR correction to account for multiple comparisons.

## Results

### Participants and Clinical Characteristics

This study included data from 493 older participants in the multi-centric DELCODE cohort (see Methods for exclusion criteria), including 203 CNs, 204 SCDs, 65 MCIs, and 21 DATs, with the group being 53.35% female and having an average age of 69.76 ± 5.65 years. A subset of 235 individuals, comprising 92 CNs, 95 SCDs, 34 MCIs, and 14 DATs (49.36% female, average age 69.70 ± 5.37 years), had complete CSF biomarker data available for subsequent brain-phenotype analyses. Demographic and clinical diagnostic characteristics, including biomarker distributions, are summarised in Supplementary Table 1.

### Dynamic Causal Model of Memory Encoding

To explore the influence of ATN pathology on memory-related EC in aging and AD, we applied DCM to fMRI data obtained during visual memory encoding, focusing on a network comprising the PPA, HC, and PCU (Figure 2B, see also methods). Individual level DCMs of memory encoding were estimated for all participants. Figure 2C illustrates a group-level connectivity profile seen in CN individuals. For task fMRI intrinsic connectivity, connections from PPA to PCU and HC to PPA were found to be inhibitory, while all other intrinsic connections were excitatory, replicating previous results in independent cohorts^28^. Our model allowed for modulation of inhibitory self-connectivity by encoding success. Successful encoding elicited a significantly negative modulatory influence on self-connectivity of both the PPA and the HC, most likely reflecting disinhibition, as evident from increased PPA and HC activation to remembered vs. forgotten items in standard GLM analysis (Figure 2a; see also Soch et al.^27^ and Vockert et al.^26^). No modulation of PCU activity related to encoding success was observed (p>0.05). The driving input to PPA was uniformly excitatory, supporting the plausibility of our model. A detailed evaluation of DCM connectivity is provided in Supplementary Table 2.

### Input-Gated Connectivity Decline is Linked to Memory Performance

We next explored a potential relationship between individual connection strengths and memory performance as defined by the AUC (A’) of the delayed recognition task (performed 90 min after the fMRI experiment; see methods) Using Spearman’s rank correlation, we identified six connectivity parameters, all involving the DCM input region PPA, whose strengths diminished significantly with decreasing memory performance (Figure 3A, supplementary table 3). Specifically, excitatory connectivity from the PPA to the HC correlated positively with A’ (ρ = 0.22, p = 2.26 × 10^⁻6^; Figure 3). Conversely, inhibitory connectivity from the PPA to the PCU correlated negatively (ρ = -0.25, p = 5.70 × 10^⁻8^), indicating poorer memory with reduced inhibitory control. Similarly, reduced inhibitory connectivity from the HC to the PPA associated with lower memory performance (ρ = -0.24, p = 2.06 × 10^⁻7^). Excitatory connectivity from the PCU to the PPA positively predicted memory performance (ρ = 0.26, p = 1.08 × 10^⁻8^), highlighting its integrative role within the memory network.

**Figure 3.**
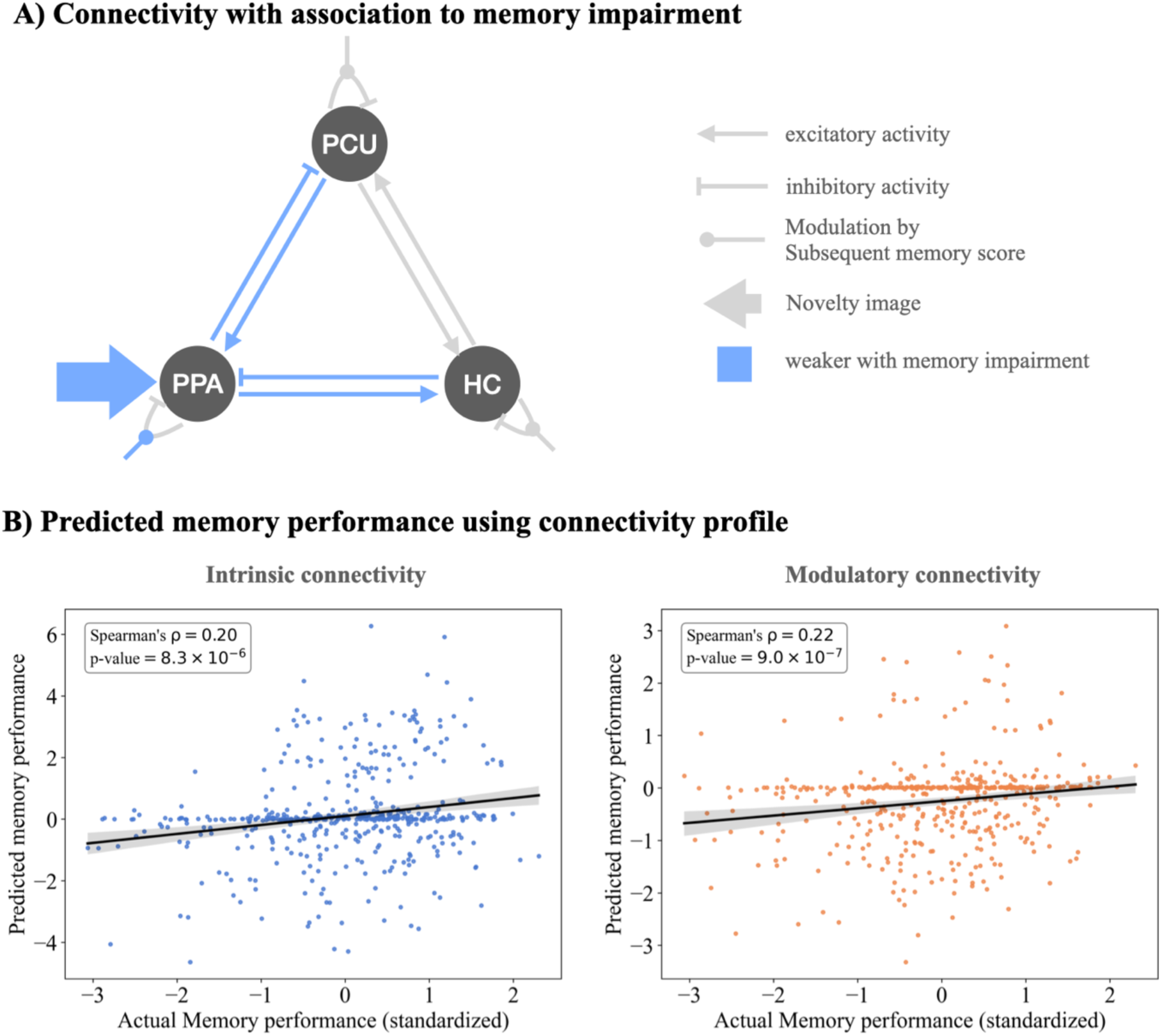
Reduced input-gated effective connectivity associates with impaired memory performance. **(A)** The connectivity strength of interregional connections, modulatory units, and external inputs related to the PPA is reduced in association with lower memory performance (see also Supplementary Table 3). These connectivity strengths are reduced in a uniform direction in association with decreased memory impairment, irrespective of whether the connectivity is excitatory or inhibitory. Significant relationships, derived from Spearman’s rank correlations between effective connectivity estimates and memory performance, are defined by *p* < 0.05 (FDR corrected) and |*ρ*| > 0.2. **(B)** The association of predicted memory performance using connectivity profiles in out-of-sample cross-validation. Left: Using the entire intrinsic connectivity profile (DCM’s A parameters), a significant positive correlation (ρ = 0.20, p = 8.3 x 10^-6^) indicates an association between memory performance and intrinsic connectivity dynamics. Right: Using the modulatory connectivity profile (DCM’s B parameters), modulatory connectivity significantly predicts memory performance (ρ = 0.22, p = 9.0 x 10^-7^), further indicating a meaningful association. Bayesian inference contributes to observed shrinkage towards zero in some subjects’ predictions. The results were derived from Spearman’s rank correlations. Abbreviations: PPA (parahippocampal place area), HC (hippocampus), PCU (precuneus).

Additionally, inhibitory self-connectivity of the PPA negatively correlated with A’ (ρ = -0.27, p = 6.14 × 10^⁻9^). Along the same line, the driving input to the PPA exhibited the strongest positive correlation with memory performance (ρ = 0.32, p = 2.66 × 10^⁻12^), both correlations likely reflecting the reduced PPA activation paralleling lower memory performance in older age^28,41^ and pre-clinical AD^26,27,49^.

Next, we aimed to validate these findings using PACC5 as an independent cognitive performance measure that is sensitive to preclinical change in Alzheimer’s disease. The correlations between connectivity and PACC5 aligned with memory performance (supplementary table 3). Excitatory connectivity from the PPA to the HC and from the PCU to the PPA showed a positive correlation with PACC5 (ρ = 0.21, p = 1.14 × 10⁻⁵ for both). Conversely, inhibitory connectivity from the PPA to the PCU and from the HC to the PPA correlated negatively with PACC5 (ρ = -0.23, p = 2.31 × 10⁻⁶ and ρ = -0.21, p = 1.14 × 10⁻⁵, respectively). The results also aligned with inhibitory self-connectivity within the PPA (ρ = - 0.24, p = 3.34 × 10⁻⁷) and the driving input to the PPA (ρ = 0.31, p = 3.66 × 10⁻¹¹).

Finally, out-of-sample cross-validation was performed in order to examine the association between connectivity profiles and cognitive performance (Figure 3B). Connectivity profiles were predictive of both memory performance and PACC5 scores. The correlations between actual and estimated memory performance were significantly positive for intrinsic connectivity (ρ = 0.20, p = 8.3 x 10^-6^) and modulatory connectivity (ρ = 0.22, p = 9.0 x 10^-7^) The same pattern was observed for PACC5, albeit with weaker associations (ρ = 0.14, p = 0.003 for intrinsic connectivity; ρ = 0.19, p = 2.3 x 10^-5^ for modulatory connectivity).

### Disruption of Forward Input-gated Connectivity is Tied to Tau and Amyloid Pathology

To assess how AD pathology might contribute to connectivity disruptions, we examined the effects of amyloid and tau biomarkers on connections associated with memory performance, accounting for potential non-linear effects and interactions (see GAMs in methods). We first focused on effects of CSF Aβ42/40 ratio and p-tau-181, which showed an association with the PPA to HC and the PPA to PCU connections (Table 1). The excitatory connection from the PPA to the HC (Figure 4A) was negatively associated with p-tau-181 levels (*p* = 0.007), reflecting a weakened excitatory PPA to HC connection in the presence of tau pathology (Figure 4B).

**Figure 4.**
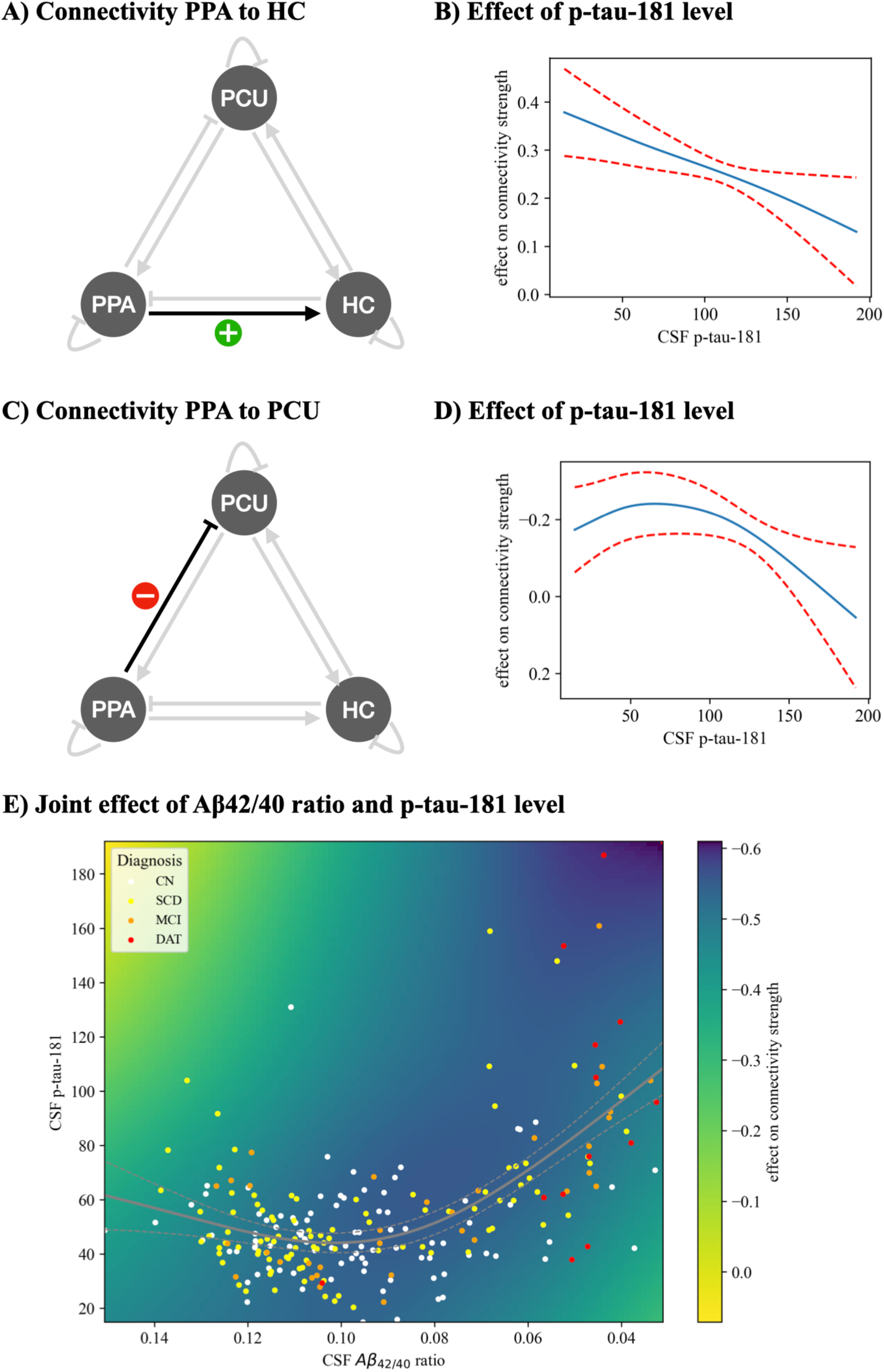
Effects of CSF amyloid-β42/40 (Aβ42/40 ratio) and CSF phospho-tau181 (p-tau-181) on forward input-gated connectivity during memory encoding. **(A)** Intrinsic connectivity from the PPA to the HC is excitatory, indicated in black. **(B)** Increasing p-tau levels are associated with a progressive reduction in the strength of excitatory connectivity from the PPA to the HC. The blue line represents effect of p-tau-181 levels, and red dashed lines denote the 95% confidence interval (CI). **(C)** Connectivity from the PPA to the PCU is inhibitory, highlighted in black. **(D)** Higher p-tau-181 levels are associated with decreased inhibitory connectivity from the PPA to the PCU. Visualization follows the same style as in (B). The y-axis is reversed to represent the effect on inhibitory connectivity (negative value). **(E)** The heatmap shows effect on the connectivity strength from the PPA to the PCU given Aβ42/40 ratio and p-tau-181 levels. The grey line represents simultaneous change of Aβ42/40 ratio and p-tau-181 levels approximated from the dataset. Toward disease progression, Aβ42/40 ratio decreases and p-tau-181 levels progressively increase, reflecting an interdependent progression. The inhibitory connectivity is decreased along with the progression. The x-axis is reversed to represent disease progression from early to late stages. The colour bar is reversed to represent the effect on inhibitory connectivity (negative value). The progression of p-tau relative to Aβ42/40 is visualised with the 95% CI shown as dashed lines. Abbreviations: PPA (parahippocampal place area), HC (hippocampus), PCU (precuneus), CN (cognitively normal), SCD (subjective cognitive decline), MCI (mild cognitive impairment), and DAT (dementia of Alzheimer’s type).

**Table 1.**
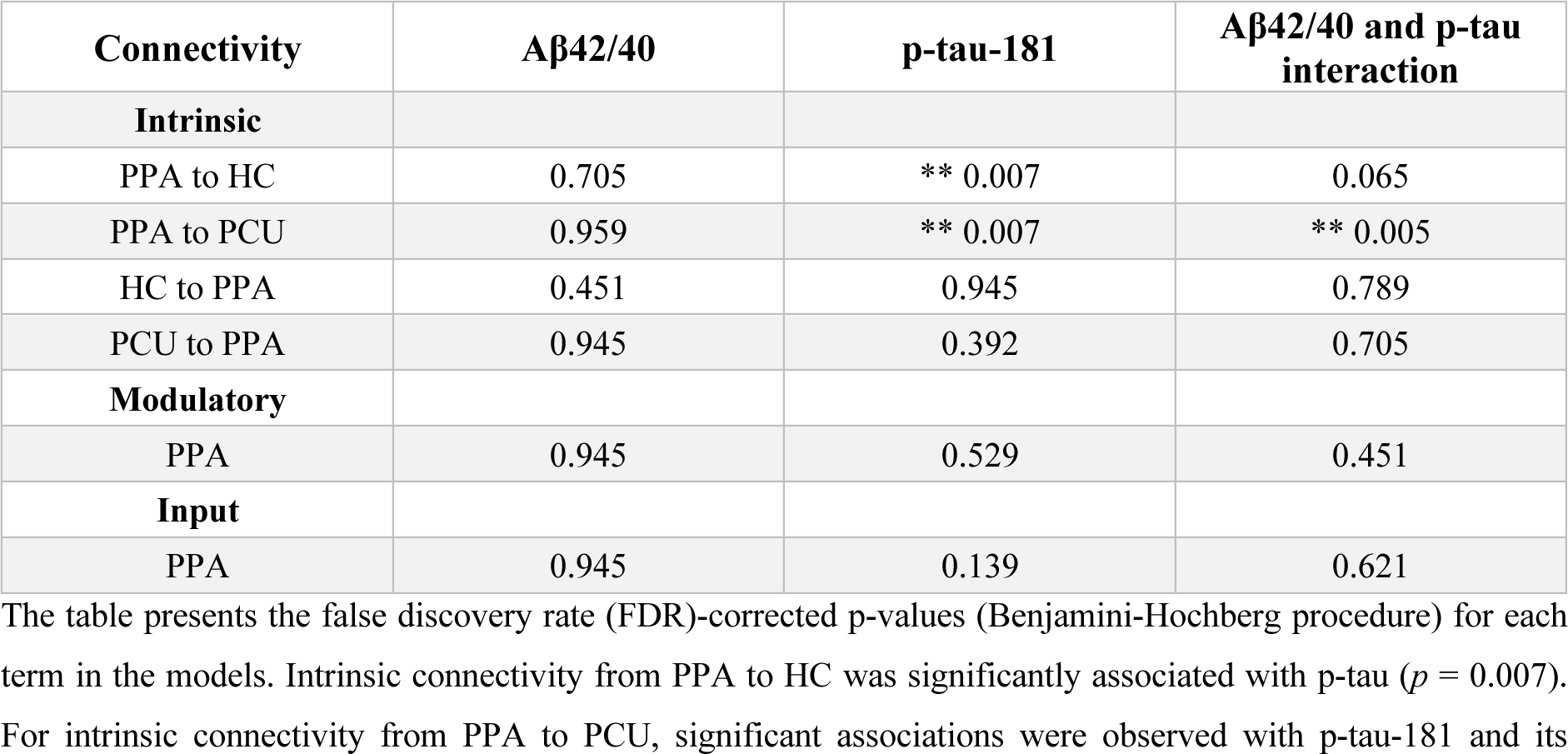
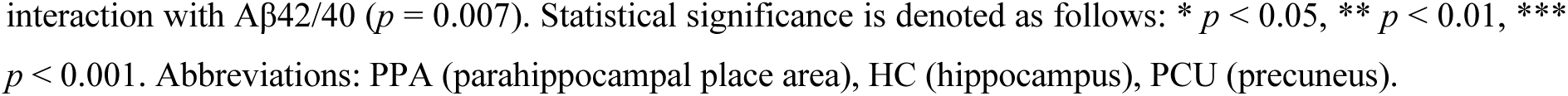
Effective connectivity during memory encoding in relation to AD pathology in terms of CSF biomarkers amyloid-β42/40 (Aβ42/40) and phospho-tau181 (p-tau) as assessed with generalised additive modelling.

| Connectivity | Aβ42/40 | p-tau-181 | Aβ42/40 and p-tau interaction |
| --- | --- | --- | --- |
| **Intrinsic** |  |  |  |
| PPA to HC | 0.705 | ** 0.007 | 0.065 |
| PPA to PCU | 0.959 | ** 0.007 | ** 0.005 |
| HC to PPA | 0.451 | 0.945 | 0.789 |
| PCU to PPA | 0.945 | 0.392 | 0.705 |
| **Modulatory** |  |  |  |
| PPA | 0.945 | 0.529 | 0.451 |
| **Input** |  |  |  |
| PPA | 0.945 | 0.139 | 0.621 |

The table presents the false discovery rate (FDR)-corrected p-values (Benjamini-Hochberg procedure) for each term in the models. Intrinsic connectivity from PPA to HC was significantly associated with p-tau ( $p = 0.007$ ). For intrinsic connectivity from PPA to PCU, significant associations were observed with p-tau-181 and its
interaction with Aβ42/40 ( $p = 0.007$ ). Statistical significance is denoted as follows: \* $p < 0.05$ , \*\* $p < 0.01$ , \*\*\* $p < 0.001$ . Abbreviations: PPA (parahippocampal place area), HC (hippocampus), PCU (precuneus).

On the other hand, the inhibitory connectivity from PPA to PCU (Figure 4C) was related to p-tau-181 levels and their interaction with the Aβ42/40 ratio. Specifically, the higher p-tau level was associated with reduced inhibitory connection strength (Figure 4D). Additionally, there was a significant interaction between p-tau-181 level and Aβ42/40 ratio on the PPA to PCU connection. To illustrate this interaction, we approximated the path of simultaneous changes of p-tau-181 level and Aβ42/40 ratio throughout the AD spectrum using spline regression (Figure 4E). This revealed that p-tau-181 levels increased with decreasing Aβ42/40 ratio. The interdependent progression of Aβ and tau pathology thus led to an amplified disruption of inhibitory connectivity. In addition to examining the effects of Aβ42/40 and p-tau, we assessed the impact of neurodegeneration—approximated by hippocampal atrophy—on the same connections of interest. Unlike Aβ42/40 and p-tau, which showed significant associations with specific connections, hippocampal volume did not exhibit a significant effect on any connection investigated (Supplementary Table 4).

### Synthesising fMRI BOLD Activity Reveals Spatially Specific Effects of Alzheimer’s Disease Pathology

After establishing that Aβ42/40 ratio and p-tau-181 levels, but not hippocampal volumes, were predictive of the EC patterns during memory encoding, we next estimated four connectivity profiles based on Aβ42/40 ratio and p-tau-181 levels for four AT pathology groups: A-T-, A+T-, A-T+, and A+T+. Four connectivity profiles corresponding to the AD pathology groups were used to synthesize BOLD signals from individual’s PPA, HC, and PCU during performing the memory-encoding task (see methods).

Synthetic BOLD for a healthy control is illustrated in Figure 5A. Following successful memory encoding, lagging positive responses were observed in PPA and HC, while negative responses were seen in PCU. These responses varied across AT classifications, demonstrating responsiveness to memory encoding associated with amyloid and tau pathology.

**Figure 5.**
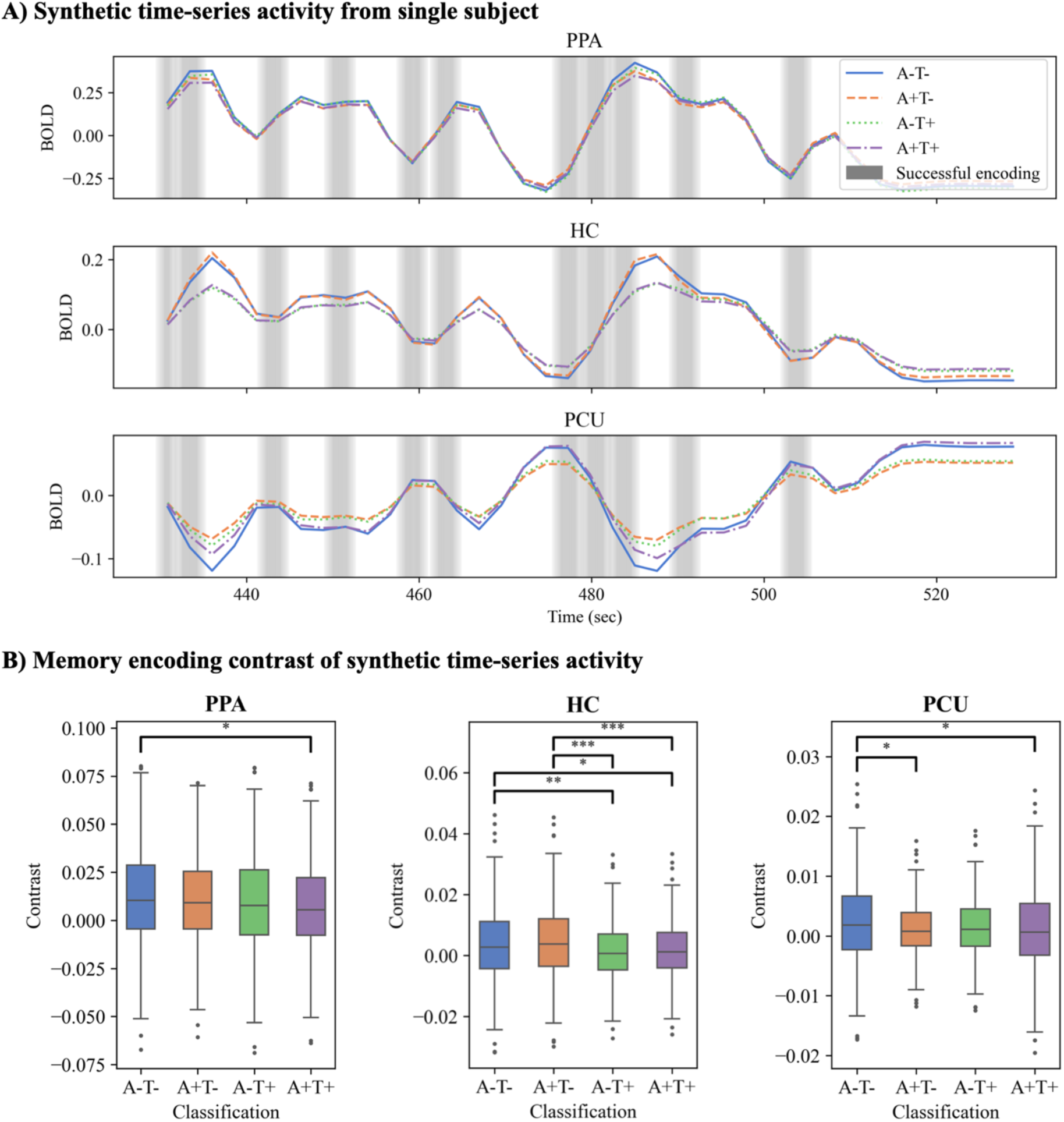
Synthesised brain activity and the comparison. **(A)** An example of synthesised activity from the region of interest (ROI). The plots demonstrate lagging positive responses in PPA and HC, and negative responses in PCU following successful encoding. **(B)** Pairwise comparisons of group-level memory contrast from the same individuals with different AD classifications show that the effects of AD pathology differ in each ROI. PPA is only affected by the simultaneous transition of amyloid (A) and tau status (T). HC is significantly affected by p-tau as reflected in the significant in the transition T- to T+. PCU is affected solely by the transition of A. Statistical significance was evaluated with Wilcoxon-rank test with False discovery rate (FDR) and is denoted as follows: * *p* < 0.05, ** *p* < 0.01, *** *p* < 0.001. Abbreviations: PPA (parahippocampal place area), HC (hippocampus), PCU (precuneus)

**Figure 6.**
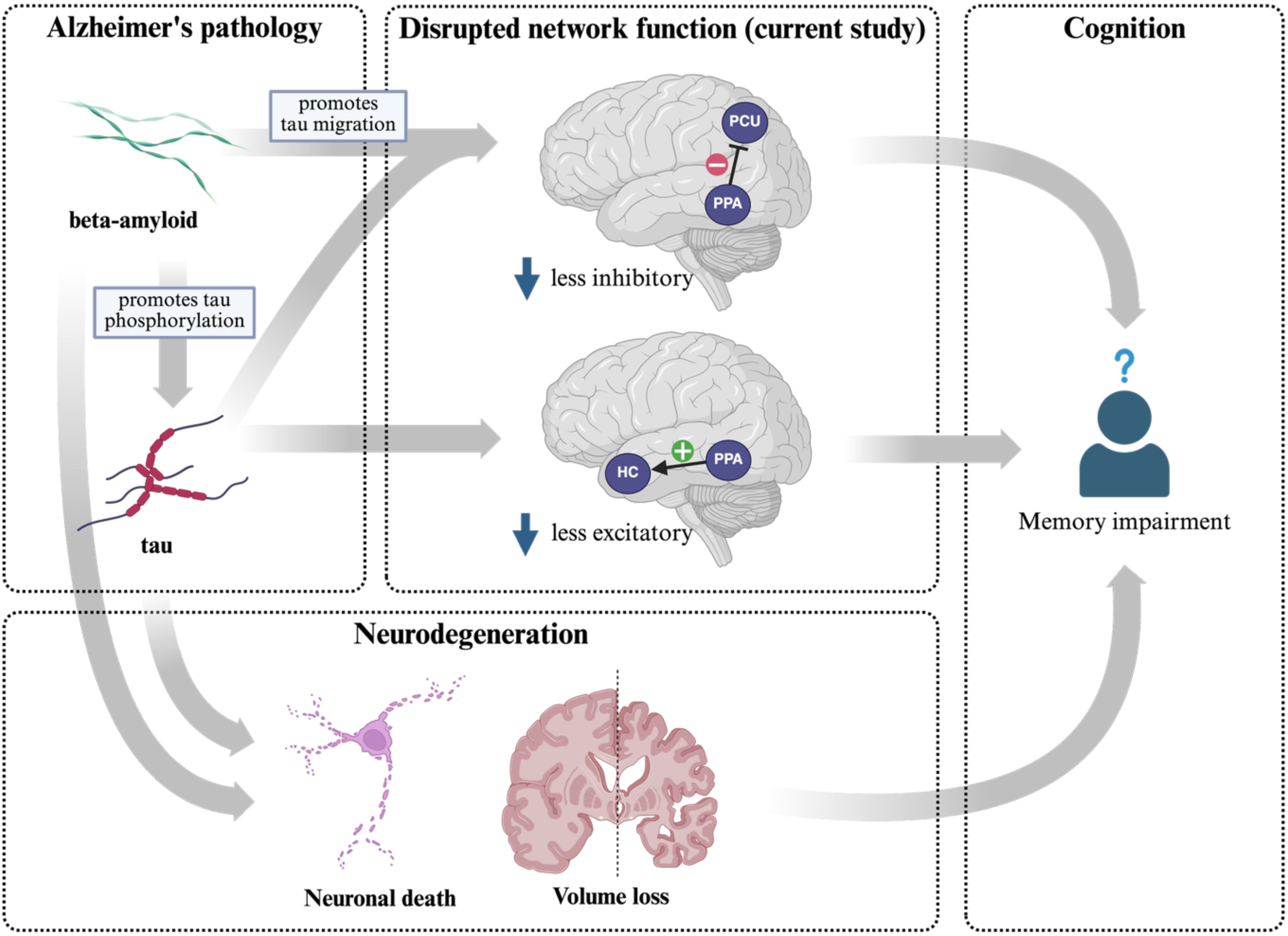
Network alterations in forward input-gated connectivity as a mechanism of cognitive impairment in Alzheimer’s disease. This illustration highlights the roles of aberrant synaptic connectivity and neurodegeneration in Alzheimer’s disease. The accumulation of beta-amyloid and p-tau proteins, central to the pathology, initiates a cascade of events. Beta-amyloid promotes tau phosphorylation and its migration from subcortical regions to the neocortex, where phosphorylated tau reduces excitatory connectivity between the hippocampal place area (PPA) and the hippocampus (HC). Simultaneously, beta-amyloid facilitates p-tau-induced decreases in inhibitory activity from the PPA to the precuneus (PCU). These synaptic aberrations, combined with synaptic and volumetric loss, ultimately lead to memory impairment.

The memory contrast revealed differential effects of Aβ42/40 and p-tau on each ROI, as shown in Figure 5B. In PPA, a significant decrease in ROI activity was observed only in the transition from A-T- to A+T+ (p = 0.013). In HC, transitions involving a change from T- to T+ consistently resulted in decreased ROI activity. Significant transitions included A-T- to A-T+ (p = 0.005), A-T- to A+T+ (p = 0.025), A+T- to A-T+ (p = 1.0 x 10^-4^), and A+T- to A+T+ (p = 5.0 x 10^-4^). For PCU, the transition from A- to A+ was associated with a reduction in ROI contrast, reflect a loss of deactivation during successful encoding. Significant changes were observed in the transitions from A-T- to A+T- (p = 0.043) and from A-T- to A+T+ (p = 0.043).

In summary, these findings from generative modelling illustrate the distinct effects of Aβ42/40 and p-tau on regional BOLD under memory encoding. PPA activity was sensitive to simultaneous changes of amyloid and tau, while HC activity was predominantly influenced by tau transitions. In contrast, PCU responses were primarily affected by amyloid changes. These contrast from synthesised data demonstrated the complexities of amyloid and tau pathology in shaping the functional dynamics of memory-related regions.

## Discussion

This study employed DCM to investigate the network dynamics underlying impaired memory-encoding process across the AD spectrum, aiming to elucidate potential mechanisms mediating the relationship between molecular biomarkers and memory performance at the level of network dysfunction. We found alterations in EC strength to predict lower memory performance in older adults within the AD spectrum, and these findings highlighted that the disruptions in forward input-gated connectivity in successful memory-encoding network play a critical role in mediating memory impairment and Alzheimer’s pathology.

### Altered memory network effective connectivity related to aging and AD

Our findings replicate the previously demonstrated importance of forward and inhibitory connectivity within the parieto-temporal network, particularly from the PPA to the HC and from the PPA to the PCU, in the encoding of novel visual information.^28^ Both connections were attenuated in individuals with worse performance in the memory encoding task (Figure 3A), a finding in line with an DCM study^28^ with the same set of ROIs, which demonstrated that an age-related weakening of inhibitory connectivity from the PPA to the PCU was associated with lower memory performance in healthy older adults. More broadly, our results support the importance of preserved inhibitory connectivity for memory functioning in old age.^50,51^

In the present study, we could expand the previous findings on age-dependent alterations of memory-related EC patterns by demonstrating that they are even more pronounced in older adults within the AD spectrum. This is mirrored by recent studies^25–27^ showing attenuated PPA activity and reduced PCU deactivation with increasing severity across the AD spectrum. Notably, our results suggest that altered EC in individuals with increased AD risk likely goes beyond accelerated aging, as it could be shown to play a role in mediating the relationship between AD molecular pathology and cognitive decline.

### Regionally specific influence of amyloid and tau pathology

Amyloid and tau are well-established biomarkers of AD and have both been shown to contribute to synaptic dysfunction and neurodegeneration.^2,5,52^ Importantly, they likely show a non-linear interaction in their contribution to AD progression: While amyloid accumulation is associated with few symptoms on its own, it plays a critical role in promoting pathological tau accumulation and influences the spatial distribution of tau by facilitating its migration from subcortical to cortical regions.^3,7,16,53–60^ On the other hand, tau appears to be more directly involved in the disruption of brain activity and cognition.^33,61–63^

Previous PET studies have demonstrated distinct anatomical patterns of amyloid and tau deposition.^64–68^ Specifically Aβ deposition in AD typically begins in the posterior cingulate cortex and connected parietal cortices^65,69^, while tau pathology originates in the medial temporal lobe (MTL), including the entorhinal and parahippocampal cortex^70^. This dissociation is, to some extent, mirrored by the observed differential influence of tau and amyloid pathology on the parieto-temporal network. Specifically, decreased excitatory activity from the PPA to the HC was linked to elevated p-tau levels, independently of amyloid, in line with previous studies^61,71^. These studies have shown that accumulation of hyperphosphorylated tau predominantly accumulates in MTL structures, particularly in parahippocampal gyrus, exhibit the strongest association with episodic memory decline. Furthermore, MTL tau level is the most reliable predictor of episodic memory performance, independent of amyloid status, but strongly linked to aging.^61^ On the other hand, decreased inhibitory activity from the PPA to the PCU was also associated with p-tau-181 levels, but with a synergistic effect of the Aβ42/40 ratio, and these were also associated with poorer memory performance. Notably, amyloid accumulation, unlike tau deposition, starts in neocortical areas, with the PCU typically being affected early in the course of AD.^65^ Compatibly, reduced deactivation of the PCU has been linked to pre-clinical AD stages^25^, and fMRI scores based on reduced activations as well as deactivations have been associated with tau and amyloid pathology alike, particularly in individuals with SCD^27^.

Taken together, our findings—namely, that connectivity from the PPA to the HC was associated with p-tau-181, while connectivity from the PPA to the PCU was influenced by p-tau-181 and its interaction with the Aβ42/40 ratio—align with these observations. They underscore the critical role of tau in altering MTL connectivity, as well as the combined impact of tau and amyloid in disrupting inhibitory interactions between the MTL and parietal or default mode network structures.

### Implications for the ATN classification of AD progression

These findings underscore the selective vulnerability of network interactions to tau pathology, particularly in the forward connectivity from the PPA to regions involved in higher-order functions. In contrast to tau, amyloid pathology did not exhibit effects on any connectivity of interest, but instead exerted influence on the inhibitory connectivity to the PCU. Moreover, the decline in forward connectivity within the parieto-temporal network along the disease trajectory was associated with cognitive impairment, as detailed in the previous section (Figure 3A). Together, the results suggest a mechanistic explanation in which tau and amyloid pathology contribute to cognitive function through the disruption of input-gated forward connectivity within the memory network

The study found no evidence that neurodegeneration, at least as far as captured by hippocampal atrophy, influenced the EC of the memory-encoding network. This result may be explained by the nonspecific nature of neurodegeneration with respect to Alzheimer’s pathology. Neurodegeneration is used as an indicator of disease severity or as a determinant of whether clinical presentation is linked to Alzheimer’s pathology.^23^ Importantly, neurodegeneration has been shown to affect cognition, including episodic memory, independently of amyloid and tau pathology.^72^ However, neurodegeneration in Alzheimer’s disease is driven both directly and indirectly by the accumulation of amyloid and tau.^53,73,74^ These findings suggest that neurodegeneration may act as an intermediary between Alzheimer’s pathology and memory impairment, potentially operating in parallel to network dysfunction. As argued by Mooraj et al.^75^ in context of aging, structural measurements may explain variance in cognitive outcomes differently than functional measurements, as they are putatively unable to capture dynamic cognitive processes and synaptic dysfunction^33^.

We leveraged the generative properties of our models to synthesise brain activity from regions of interest for the same individuals under different AT classifications. This approach allowed us to discern the effects of amyloid and tau pathology while controlling for individual-specific factors and experimental conditions. Moreover, it helped mitigate the issue of limited statistical power, particularly for the relatively uncommon A-T+ classification (see Heinzinger et al.^34^ which used the same dataset; see Walsh et al.^76^ that proposed similar idea for randomised clinical trial; see Ritter et al.^77^ for a greater scale of brain functional simulation). Our findings highlighted the susceptibility of the hippocampal activity to the AD pathology, which was specifically affected by various transitions involving an increasing tau (T) status. In contrast, the activity of the PCU was primarily sensitive to changes in amyloid (A) status. This distinction underscores the spatial specificity and differential vulnerability of brain regions in the progression of AD, where tau tangles early accumulate in the MTL and amyloid-beta plaques in the precuneus.^65,78^ Furthermore, in assessing the individual effects of amyloid and tau pathology, our synthetic data successfully reproduced findings from previous studies^33,65^, reinforcing the robustness and validity of our approach.

We present a mechanistic model linking molecular pathology to memory impairment in AD, emphasizing network disruptions as a critical pathological intermediary (Figure 5). Tau is identified as the primary agent driving connectivity aberrations, with amyloid exerting a synergistic effect by promoting tau phosphorylation and indirectly facilitating tau migration from subcortical to cortical regions. This process leads to decreased excitatory connectivity within the MTL (connectivity from PPA to HC) and decreased inhibitory connectivity between the MTL and the posteromedial cortex (connectivity from PPA to PCU), ultimately resulting in memory impairment. The absence of a direct effect of neurodegeneration on connectivity suggests a parallel pathological pathway in which amyloid and tau together contribute to both synaptic and neuronal loss, culminating in cognitive decline.

### Limitations

This study has several limitations. First, the proportion of participants in the more severe stages of the AD spectrum was relatively low, which may have limited the ability to detect biomarker effects in the terminal stages of the disease. This limitation was further compounded by the challenges faced by participants with dementia in complying with task-based fMRI protocols, resulting in their exclusion from the study. Second, we were unable to extract time-series data using a threshold for significant voxel activity. This limitation arose because the absence of significant voxels was associated with reduced hippocampal volume and poorer memory performance. As recently shown, individuals in the later stages of the AD spectrum exhibit diminished fMRI subsequent memory effects^79^, which would have resulted in a systematic exclusion of individuals at later stages. Aware of the trade-off between data quality and inclusivity, we chose to prioritise a comprehensive representation of the AD spectrum, ensuring the characteristics of the late stages were captured. Third, the CSF biomarkers used in this study lacked spatial specificity, which is informative for understanding complex interactions involving regionally specific aggregation and protein migration. Finally, there are several concerns regarding DCM, including a lack of independent validation and potential oversimplification.^80,81^ To address these issues, we implemented several procedures, such as adopting theory-driven models and enhancing transparency through cross-validation methods.

## Conclusions

In conclusion, this study provides critical insights into the mechanisms by which Alzheimer’s pathology impairs memory performance, emphasizing the importance of disrupted forward connectivity within the parieto-temporal network during memory encoding. Specifically, the findings highlight that decreased excitatory connectivity from PPA to HC, driven by elevated p-tau levels, and reduced inhibitory connectivity from PPA to PCU, modulated by the interaction between amyloid and tau, play pivotal roles in memory impairment. These results advance our understanding of how molecular pathology translates to network-level dysfunction and cognitive decline. By identifying specific connectivity pathways linked to Alzheimer’s biomarkers, this work opens new avenues for targeted interventions aimed at mitigating memory deficits through the preservation or restoration of synaptic function.

## Data availability

Data, study protocol, and biomaterials can be shared with partners based on individual data and biomaterial transfer agreements. Access to the relevant study data can be obtained by submitting an application to the Clinical Research Platform of the DZNE (https://www.dzne.de/en/research/research-areas/clinical-research/for-researchers/).

## Acknowledgements

Not applicable.

## Abbreviations

AB: beta-amyloid
AD: Alzheimer’s disease
CN: cognitively normal
DAT: dementia of Alzheimer’s type
DCM: Dynamic causal modelling
DELCODE: DZNE - Longitudinal Cognitive Impairment and Dementia Study
DZNE: German Centre for Neurodegenerative Diseases
GAMs: Generalised additive models
HC: hippocampus
MCI: mild cognitive impairment
PPA: parahippocampal place area
PCU: precuneus
p-tau: phosphorylated tau
SCD: subjective cognitive decline

## References

1. Jack CR, Knopman DS, Jagust WJ, et al. Tracking pathophysiological processes in Alzheimer’s disease: an updated hypothetical model of dynamic biomarkers. The Lancet Neurology. 2013;12(2):207–216. doi:10.1016/s1474-4422(12)70291-0

2. Tzioras M, McGeachan RI, Durrant CS, Spires-Jones TL. Synaptic degeneration in Alzheimer disease. Nature Reviews Neurology. 2023;19(1):19–38. doi:10.1038/s41582-022-00749-z

3. Busche MA, Hyman BT. Synergy between amyloid-β and tau in Alzheimer’s disease. Nature Neuroscience. 2020;23(10):1183–1193. doi:10.1038/s41593-020-0687-6

4. Heneka MT, Carson MJ, Khoury JE, et al. Neuroinflammation in Alzheimer’s disease. The Lancet Neurology. 2015;14(4):388–405. doi:10.1016/s1474-4422(15)70016-5

5. Selkoe DJ. Alzheimer’s disease is a synaptic failure. Science. 2002;298(5594):789-791. doi:10.1126/science.1074069

6. Reddy PH, Beal MF. Amyloid beta, mitochondrial dysfunction and synaptic damage: implications for cognitive decline in aging and Alzheimer’s disease. Trends in Molecular Medicine. 2008;14(2):45–53. doi:10.1016/j.molmed.2007.12.002

7. Bennett RE, DeVos SL, Dujardin S, et al. Enhanced Tau Aggregation in the Presence of Amyloid β. The American Journal of Pathology. 2017;187(7):1601–1612. doi:10.1016/j.ajpath.2017.03.011

8. Goedert M, Spillantini MG, Crowther RA. Tau Proteins and Neurofibrillary Degeneration. Brain Pathology. 1991;1(4):279-286. doi:10.1111/j.1750-3639.1991.tb00671.x

9. Stokin GB, Lillo C, Falzone TL, et al. Axonopathy and Transport Deficits Early in the Pathogenesis of Alzheimer’s Disease. Science. 2005;307(5713):1282-1288. doi:10.1126/science.1105681

10. Thal DR, Rüb U, Orantes M, Braak H. Phases of Aβ-deposition in the human brain and its relevance for the development of AD. Neurology. 2002;58(12):1791–1800. doi:10.1212/wnl.58.12.1791

11. Mattsson-Carlgren N, Salvadó G, Ashton NJ, et al. Prediction of Longitudinal Cognitive Decline in Preclinical Alzheimer Disease Using Plasma Biomarkers. JAMA Neurology. 2023;80(4):360. doi:10.1001/jamaneurol.2022.5272

12. Braak H, Braak E. Neuropathological stageing of Alzheimer-related changes. Acta Neuropathologica. 1991;82(4):239–259. doi:10.1007/bf00308809

13. Braak H, Braak E. Frequency of Stages of Alzheimer-Related Lesions in Different Age Categories. Neurobiology of Aging. 1997;18(4):351–357. doi:10.1016/s0197-4580(97)00056-0

14. Hampel H, Hardy J, Blennow K, et al. The Amyloid-β Pathway in Alzheimer’s Disease. Molecular Psychiatry. 2021;26(10):5481–5503. doi:10.1038/s41380-021-01249-0

15. Roemer-Cassiano SN, Wagner F, Evangelista L, et al. Amyloid-associated hyperconnectivity drives tau spread across connected brain regions in Alzheimer’s disease. Science Translational Medicine. 17(782):eadp2564. doi:10.1126/scitranslmed.adp2564

16. Vogel JW, Iturria-Medina Y, Strandberg OT, et al. Spread of pathological tau proteins through communicating neurons in human Alzheimer’s disease. Nature Communications. 2020;11(1):2612. doi:10.1038/s41467-020-15701-2

17. Giorgio J, Adams JN, Maass A, Jagust WJ, Breakspear M. Amyloid induced hyperexcitability in default mode network drives medial temporal hyperactivity and early tau accumulation. Neuron. Published online 2023. doi:10.1016/j.neuron.2023.11.014

18. Friston KJ. Functional and Effective Connectivity: A Review. Brain Connectivity. 2011;1(1):13–36. doi:10.1089/brain.2011.0008

19. Friston KJ. Functional and effective connectivity in neuroimaging: A synthesis. Human Brain Mapping. 1994;2(1):56–78. doi:10.1002/hbm.460020107

20. Friston KJ, Harrison L, Penny W. Dynamic causal modelling. NeuroImage. 2003;19(4):1273-1302. doi:10.1016/s1053-8119(03)00202-7

21. Zeidman P, Jafarian A, Corbin N, et al. A guide to group effective connectivity analysis, part 1: First level analysis with DCM for fMRI. NeuroImage. 2019;200:174–190. doi:10.1016/j.neuroimage.2019.06.031

22. Jack CR, Bennett DA, Blennow K, et al. A/T/N: An unbiased descriptive classification scheme for Alzheimer disease biomarkers. Neurology. 2016;87(5):539–547. doi:10.1212/wnl.0000000000002923

23. Jack CR, Bennett DA, Blennow K, et al. NIA-AA Research Framework: Toward a biological definition of Alzheimer’s disease. Alzheimer’s & dementia : the journal of the Alzheimer’s Association. 2018;14(4):535–562. doi:10.1016/j.jalz.2018.02.018

24. Sperling RA, Dickerson BC, Pihlajamaki M, et al. Functional Alterations in Memory Networks in Early Alzheimer’s Disease. Neuromol Med. 2010;12(1):27–43. doi:10.1007/s12017-009-8109-7

25. Billette OV, Ziegler G, Aruci M, et al. Novelty-Related fMRI Responses of Precuneus and Medial Temporal Regions in Individuals at Risk for Alzheimer Disease. Neurology. 2022;99(8):e775–e788. doi:10.1212/wnl.0000000000200667

26. Vockert N, Machts J, Kleineidam L, et al. Cognitive reserve against Alzheimer’s pathology is linked to brain activity during memory formation. Nature Communications. 2024;15(1):9815. doi:10.1038/s41467-024-53360-9

27. Soch J, Richter A, Kizilirmak JM, et al. Single-value brain activity scores reflect both severity and risk across the Alzheimer’s continuum. Brain. 2024;147(11):3789–3803. doi:10.1093/brain/awae149

28. Schott BH, Soch J, Kizilirmak JM, et al. Inhibitory temporo-parietal effective connectivity is associated with explicit memory performance in older adults. iScience. 2023;26(10):107765. doi:10.1016/j.isci.2023.107765

29. Jessen F, Spottke A, Boecker H, et al. Design and first baseline data of the DZNE multicenter observational study on predementia Alzheimer’s disease (DELCODE). Alzheimer’s Research & Therapy. 2018;10(1):15. doi:10.1186/s13195-017-0314-2

30. Hinrichs H, Scholz M, Tempelmann C, Woldorff MG, Dale AM, Heinze HJ. Deconvolution of Event-Related fMRI Responses in Fast-Rate Experimental Designs: Tracking Amplitude Variations. Journal of Cognitive Neuroscience. 2000;12(Supplement 2):76–89. doi:10.1162/089892900564082

31. Soch J, Richter A, Schütze H, et al. Bayesian model selection favors parametric over categorical fMRI subsequent memory models in young and older adults. NeuroImage. 2021;230:117820. doi:10.1016/j.neuroimage.2021.117820

32. Soch J, Richter A, Schütze H, et al. A comprehensive score reflecting memory-related fMRI activations and deactivations as potential biomarker for neurocognitive aging. Human Brain Mapping. 2021;42(14):4478–4496. doi:10.1002/hbm.25559

33. Düzel E, Ziegler G, Berron D, et al. Amyloid pathology but not APOE ε4 status is permissive for tau-related hippocampal dysfunction. Brain. 2022;145(4):1473–1485. doi:10.1093/brain/awab405

34. Heinzinger N, Maass A, Berron D, et al. Exploring the ATN classification system using brain morphology. Alzheimer’s Research & Therapy. 2023;15(1):50. doi:10.1186/s13195-023-01185-x

35. Papp KV, Rentz DM, Orlovsky I, Sperling RA, Mormino EC. Optimizing the preclinical Alzheimer’s cognitive composite with semantic processing: The PACC5. Alzheimer’s & Dementia: Translational Research & Clinical Interventions. 2017;3(4):668–677. doi:10.1016/j.trci.2017.10.004

36. Papp KV, Buckley R, Mormino E, et al. Clinical meaningfulness of subtle cognitive decline on longitudinal testing in preclinical AD. Alzheimer’s & Dementia. 2020;16(3):552–560. doi:10.1016/j.jalz.2019.09.074

37. Tzourio-Mazoyer N, Landeau B, Papathanassiou D, et al. Automated Anatomical Labeling of Activations in SPM Using a Macroscopic Anatomical Parcellation of the MNI MRI Single-Subject Brain. NeuroImage. 2002;15(1):273–289. doi:10.1006/nimg.2001.0978

38. Köhler S, Crane J, Milner B. Differential contributions of the parahippocampal place area and the anterior hippocampus to human memory for scenes. Hippocampus. 2002;12(6):718–723. doi:10.1002/hipo.10077

39. Kremers NAW, Deuker L, Kranz TA, Oehrn C, Fell J, Axmacher N. Hippocampal control of repetition effects for associative stimuli. Hippocampus. 2014;24(7):892–902. doi:10.1002/hipo.22278

40. Dennis NA, Hayes SM, Prince SE, Madden DJ, Huettel SA, Cabeza R. Effects of Aging on the Neural Correlates of Successful Item and Source Memory Encoding. *Journal of Experimental Psychology: Learning*, Memory, and Cognition. 2008;34(4):791–808. doi:10.1037/0278-7393.34.4.791

41. Maillet D, Rajah MN. Age-related differences in brain activity in the subsequent memory paradigm: A meta-analysis. Neuroscience & Biobehavioral Reviews. 2014;45:246–257. doi:10.1016/j.neubiorev.2014.06.006

42. Kizilirmak JM, Soch J, Schütze H, et al. The relationship between resting-state amplitude fluctuations and memory-related deactivations of the default mode network in young and older adults. Human Brain Mapping. 2023;44(9):3586–3609. doi:10.1002/hbm.26299

43. Kim H. Neural activity that predicts subsequent memory and forgetting: A meta-analysis of 74 fMRI studies. NeuroImage. 2011;54(3):2446–2461. doi:10.1016/j.neuroimage.2010.09.045

44. Friston K, Mattout J, Trujillo-Barreto N, Ashburner J, Penny W. Variational free energy and the Laplace approximation. NeuroImage. 2007;34(1):220–234. doi:10.1016/j.neuroimage.2006.08.035

45. Zeidman P, Friston K, Parr T. A Primer on Variational Laplace (VL). NeuroImage. Published online August 2023:120310. doi:10.1016/j.neuroimage.2023.120310

46. Benjamini Y, Hochberg Y. Controlling the False Discovery Rate: A Practical and Powerful Approach to Multiple Testing. Journal of the Royal Statistical Society Series B: Statistical Methodology. 1995;57(1):289–300. doi:10.1111/j.2517-6161.1995.tb02031.x

47. Zeidman P, Jafarian A, Seghier ML, et al. A guide to group effective connectivity analysis, part 2: Second level analysis with PEB. NeuroImage. 2019;200:12–25. doi:10.1016/j.neuroimage.2019.06.032

48. Servén D, Brummitt C, Abedi H, Hlink. dswah/pyGAM: v0.8.0. Published online October 31, 2018. doi:10.5281/ZENODO.1476122

49. Lattmann-Grefe R, Vockert N, Machts J, et al. Dysfunction of the episodic memory network in the Alzheimer’s disease cascade. bioRxiv. Published online 2024. doi:10.1101/2024.10.25.620237

50. Nyberg L, Andersson M, Lundquist A, Salami A, Wåhlin A. Frontal Contribution to Hippocampal Hyperactivity During Memory Encoding in Aging. Frontiers in Molecular Neuroscience. 2019;12:229. doi:10.3389/fnmol.2019.00229

51. Diersch N, Valdes-Herrera JP, Tempelmann C, Wolbers T. Increased Hippocampal Excitability and Altered Learning Dynamics Mediate Cognitive Mapping Deficits in Human Aging. The Journal of Neuroscience. 2021;41(14):3204–3221. doi:10.1523/jneurosci.0528-20.2021

52. LaFerla FM, Oddo S. Alzheimer’s disease: Aβ, tau and synaptic dysfunction. Trends in Molecular Medicine. 2005;11(4):170–176. doi:10.1016/j.molmed.2005.02.009

53. Bilgel M, Wong DF, Moghekar AR, Ferrucci L, Resnick SM, Initiative the ADN. Causal links among amyloid, tau, and neurodegeneration. Brain Communications. 2022;4(4):fcac193. doi:10.1093/braincomms/fcac193

54. Schoonhoven DN, Coomans EM, Millán AP, et al. Tau protein spreads through functionally connected neurons in Alzheimer’s disease: a combined MEG/PET study. Brain. 2023;146(10):4040–4054. doi:10.1093/brain/awad189

55. Wang L, Benzinger TL, Su Y, et al. Evaluation of Tau Imaging in Staging Alzheimer Disease and Revealing Interactions Between β-Amyloid and Tauopathy. JAMA Neurology. 2016;73(9):1070. doi:10.1001/jamaneurol.2016.2078

56. Pontecorvo MJ, Sr MDD, Navitsky M, et al. Relationships between flortaucipir PET tau binding and amyloid burden, clinical diagnosis, age and cognition. Brain. 2017;140(3):748–763. doi:10.1093/brain/aww334

57. Hurtado DE, Molina-Porcel L, Iba M, et al. Aβ Accelerates the Spatiotemporal Progression of Tau Pathology and Augments Tau Amyloidosis in an Alzheimer Mouse Model. The American Journal of Pathology. 2010;177(4):1977–1988. doi:10.2353/ajpath.2010.100346

58. Jacobs HIL, Hedden T, Schultz AP, et al. Structural tract alterations predict downstream tau accumulation in amyloid-positive older individuals. Nature Neuroscience. 2018;21(3):424–431. doi:10.1038/s41593-018-0070-z

59. Wuestefeld A, Binette AP, Berron D, et al. Age-related and amyloid-beta-independent tau deposition and its downstream effects. Brain. 2023;146(8):3192–3205. doi:10.1093/brain/awad135

60. Cody KA, Langhough RE, Zammit MD, et al. Characterizing brain tau and cognitive decline along the amyloid timeline in Alzheimer’s disease. Brain. 2024;147(6):2144–2157. doi:10.1093/brain/awae116

61. Maass A, Lockhart SN, Harrison TM, et al. Entorhinal Tau Pathology, Episodic Memory Decline, and Neurodegeneration in Aging. The Journal of Neuroscience. 2018;38(3):530–543. doi:10.1523/jneurosci.2028-17.2017

62. Busche MA, Wegmann S, Dujardin S, et al. Tau impairs neural circuits, dominating amyloid-β effects, in Alzheimer models in vivo. Nature Neuroscience. 2019;22(1):57–64. doi:10.1038/s41593-018-0289-8

63. Hanseeuw BJ, Betensky RA, Jacobs HIL, et al. Association of Amyloid and Tau With Cognition in Preclinical Alzheimer Disease. JAMA Neurology. 2019;76(8):915–924. doi:10.1001/jamaneurol.2019.1424

64. Maass A, Berron D, Harrison TM, et al. Alzheimer’s pathology targets distinct memory networks in the ageing brain. Brain. 2019;142(8):2492–2509. doi:10.1093/brain/awz154

65. Palmqvist S, Schöll M, Strandberg O, et al. Earliest accumulation of β-amyloid occurs within the default-mode network and concurrently affects brain connectivity. Nature Communications. 2017;8(1):1214. doi:10.1038/s41467-017-01150-x

66. Villain N, Chételat G, Grassiot B, et al. Regional dynamics of amyloid-β deposition in healthy elderly, mild cognitive impairment and Alzheimer’s disease: a voxelwise PiB–PET longitudinal study. Brain. 2012;135(7):2126–2139. doi:10.1093/brain/aws125

67. Vemuri P, Lowe VJ, Knopman DS, et al. Tau-PET uptake: Regional variation in average SUVR and impact of amyloid deposition. Alz & Dem Diag Ass & Dis Mo. 2017;6(1):21–30. doi:10.1016/j.dadm.2016.12.010

68. Ossenkoppele R, Schonhaut DR, Schöll M, et al. Tau PET patterns mirror clinical and neuroanatomical variability in Alzheimer’s disease. Brain. 2016;139(5):1551–1567. doi:10.1093/brain/aww027

69. Buckner RL, Snyder AZ, Shannon BJ, et al. Molecular, Structural, and Functional Characterization of Alzheimer’s Disease: Evidence for a Relationship between Default Activity, Amyloid, and Memory. The Journal of Neuroscience. 2005;25(34):7709–7717. doi:10.1523/jneurosci.2177-05.2005

70. Schöll M, Lockhart SN, Schonhaut DR, et al. PET Imaging of Tau Deposition in the Aging Human Brain. Neuron. 2016;89(5):971–982. doi:10.1016/j.neuron.2016.01.028

71. Crary JF, Trojanowski JQ, Schneider JA, et al. Primary age-related tauopathy (PART): a common pathology associated with human aging. Acta Neuropathol. 2014;128(6):755–766. doi:10.1007/s00401-014-1349-0

72. Tanner JA, Iaccarino L, Edwards L, et al. Amyloid, tau and metabolic PET correlates of cognition in early and late-onset Alzheimer’s disease. Brain. 2022;145(12):4489–4505. doi:10.1093/brain/awac229

73. Marks SM, Lockhart SN, Baker SL, Jagust WJ. Tau and β-Amyloid Are Associated with Medial Temporal Lobe Structure, Function, and Memory Encoding in Normal Aging. The Journal of Neuroscience. 2017;37(12):3192–3201. doi:10.1523/jneurosci.3769-16.2017

74. Fortea J, Vilaplana E, Alcolea D, et al. Cerebrospinal fluid β-amyloid and phospho-tau biomarker interactions affecting brain structure in preclinical Alzheimer disease. Annals of Neurology. 2014;76(2):223–230. doi:10.1002/ana.24186

75. Mooraj Z, Salami A, Campbell KL, et al. Toward a functional future for the cognitive neuroscience of human aging. Neuron. 2025;113(1):154–183. doi:10.1016/j.neuron.2024.12.008

76. Walsh JR, Roumpanis S, Bertolini D, Delmar P. Evaluating Digital Twins for Alzheimer’s Disease using Data from a Completed Phase 2 Clinical Trial. Alzheimer’s & Dementia. 2022;18(S10):e065386. doi:10.1002/alz.065386

77. Ritter P, Schirner M, McIntosh AR, Jirsa VK. The Virtual Brain Integrates Computational Modeling and Multimodal Neuroimaging. Brain Connectivity. 2013;3(2):121–145. doi:10.1089/brain.2012.0120

78. Berron D, Vogel JW, Insel PS, et al. Early stages of tau pathology and its associations with functional connectivity, atrophy and memory. Brain. 2021;144(9):2771–2783. doi:10.1093/brain/awab114

79. Soch J, Richter A, Kizilirmak JM, et al. Reduced expression of fMRI subsequent memory effects with increasing severity across the Alzheimer’s disease risk spectrum. Imaging Neuroscience. 2024;2:1–23. doi:10.1162/imag_a_00260

80. Lohmann G, Erfurth K, Müller K, Turner R. Critical comments on dynamic causal modelling. NeuroImage. 2012;59(3):2322–2329. doi:10.1016/j.neuroimage.2011.09.025

81. Daunizeau J, David O, Stephan KE. Dynamic causal modelling: A critical review of the biophysical and statistical foundations. NeuroImage. 2011;58(2):312–322. doi:10.1016/j.neuroimage.2009.11.062

