## Supplementary for "Disrupted Forward Connectivity in Parieto-temporal Network Impairs Memory Performance in Alzheimer’s Disease"

### Supplementary material

#### Supplementary table 1 Characteristics of the samples in the study by analyses.

| | | | | | |
|---|---|---|---|---|---|
| **Connectivity-Memory performance correlation (n = 493)** | | | | | |
| | **CN** | **SCD** | **MCI** | **DAT** | **all** |
| n | 203 | 204 | 65 | 21 | 493 |
| age (yr) | 68.17 (5.14) | 70.07 (5.88) | 72.63 (4.76) | 73.36 (5.38) | 69.76 (5.65) |
| Sex (% female) | 61.58% | 44.12% | 52.31% | 66.67% | 53.35% |
| education (yr) | 14.55 (2.72) | 15.25 (2.90) | 13.43 (2.81) | 13.71 (2.80) | 14.66 (2.87) |
| Memory performance | 0.76 (0.08) | 0.76 (0.09) | 0.69 (0.10) | 0.60 (0.08) | 0.75 (0.09) |
| **Biomarker-Connectivity relationship (n = 235)** | | | | | |
| | **CN** | **SCD** | **MCI** | **DAT** | **all** |
| n | 92 | 95 | 34 | 14 | 235 |
| age (yr) | 67.95 (4.98) | 70.05 (5.39) | 72.16 (4.54) | 72.91 (5.81) | 69.70 (5.37) |
| Sex (% female) | 55.43% | 40.00% | 50.00% | 71.43% | 49.36% |
| education (yr) | 14.43 (2.70) | 15.31 (2.87) | 13.38 (2.31) | 13.86 (2.35) | 14.60 (2.77) |
| Memory performance | 0.76 (0.08) | 0.77 (0.09) | 0.67 (0.09) | 0.59 (0.09) | 0.74 (0.10) |
| Aβ42/40 ratio | 0.10 (0.02) | 0.10 (0.03) | 0.08 (0.03) | 0.05 (0.02) | 0.09 (0.03) |
| p-tau (pg/ml) | 48.11 (16.89) | 54.50 (23.64) | 61.68 (29.38) | 97.56 (52.52) | 55.61 (27.27) |
| Hippocampal volume | 3658.79 (421.36) | 3687.80 (440.22) | 3146.51 (516.20) | 2800.16 (564.02) | 3545.25 (521.32) |

The table provides an overview of the characteristics of the subsamples used in the study. The upper section summarises the subsample analysed with Spearman's rank correlation to assess the relationship between effective connectivity and memory performance. The lower section details the subsample examined using a Generalised Additive Model (GAM) to explore the relationship between ATN biomarkers and effective connectivity. Summary statistics are reported as means, with standard deviations in parentheses. Abbreviations: CN (cognitively normal), SCD (subjective cognitive decline), MCI (mild cognitive impairment), and DAT (dementia of Alzheimer's type).

**Supplementary table 2 Estimated effective connectivity from dynamic causal modelling from memory-encoding task.**

| Connectivity | CN (n = 203) | SCD (n = 204) | MCI (n = 65) | DAT (n = 21) | all (n=235) | Comparison |
|---|---|---|---|---|---|---|
| **Intrinsic** | | | | | | |
| PPA to PPA | 0.08(0.11)* | 0.07(0.10)* | 0.07(0.10)* | 0.03(0.09)* | 0.08(0.11)* | CN>DAT, SCD>DAT |
| PPA to HC | 0.26(0.21)* | 0.28(0.19)* | 0.24(0.18)* | 0.12(0.19)* | 0.26(0.20)* | CN>DAT, SCD>DAT, MCI>DAT |
| PPA to PCU | -0.20(0.21)* | -0.16(0.22)* | -0.09(0.21)* | -0.11(0.17)* | -0.17(0.21)* | CN<MCI, SCD<MCI |
| HC to PPA | -0.14(0.23)* | -0.16(0.24)* | -0.14(0.17)* | 0.00(0.13) | -0.14(0.22)* | CN<DAT, SCD<DAT, MCI<DAT |
| HC to HC | -0.03(0.11)* | -0.02(0.10)* | -0.03(0.08)* | -0.02(0.07) | -0.03(0.10)* | |
| HC to PCU | 0.06(0.20)* | 0.06(0.19)* | 0.06(0.17)* | 0.04(0.12)* | 0.06(0.19)* | |
| PCU to PPA | 0.20(0.25)* | 0.15(0.26)* | 0.08(0.24)* | 0.11(0.20)* | 0.16(0.25)* | CN>MCI, CN>DAT, SCD>MCI |
| PCU to HC | 0.05(0.19)* | 0.05(0.19)* | 0.00(0.15) | 0.01(0.15) | 0.04(0.19)* | |
| PCU to PCU | -0.03(0.11)* | -0.02(0.10)* | -0.02(0.06)* | -0.02(0.05) | -0.02(0.10)* | |
| **Modulation** | | | | | | |
| PPA | -0.60(0.78)* | -0.71(0.85)* | -0.30(0.61)* | -0.26(0.72) | -0.59(0.80)* | CN<MCI, CN<DAT, SCD<MCI, SCD<DAT |
| HC | -0.15(0.98)* | -0.21(0.98)* | 0.00(1.01) | -0.17(1.10) | -0.15(0.99)* | SCD<MCI |
| PCU | -0.18(1.11) | -0.19(1.00)* | -0.18(0.91) | -0.12(0.79) | -0.18(1.03)* | |
| **Input** | | | | | | |
| PPA | 0.38(0.22)* | 0.37(0.22)* | 0.30(0.20)* | 0.19(0.13)* | 0.36(0.22)* | CN>MCI, CN>DAT, SCD>MCI, SCD>DAT, MCI>DAT |

The table presents the estimated effective connectivity derived from dynamic causal modelling for each connection and diagnostic group. Summary statistics are reported as means, with standard deviations in parentheses. Connectivity parameters from each group were tested against zero with Wilcoxon sum rank test corrected for false discovery rate (FDR). ′*′ denotes $p < 0.05$. The column comparison shows the statistically different pair of diagnostic groups by each connectivity evaluated with Mann-Whitney U test corrected for FDR. Abbreviations: PPA (parahippocampal place area), HC (hippocampus), and PCU (precuneus), CN (cognitively normal), SCD (subjective cognitive decline), MCI (mild cognitive impairment), and DAT (dementia of Alzheimer's type).

**Supplementary table 3 Spearman's rank correlation of effective connectivity and memory performance.**

| Connectivity | Memory performance | | | PACC5 | | |
|---|---|---|---|---|---|---|
| | correlation | p-value | significance | correlation | p-value | significance |
| **Intrinsic** | | | | | | |
| PPA to PPA | 0.11 | 0.031 | . | 0.13 | 0.007 | . |
| PPA to HC | 0.22 | $2.26 \times 10^{-6}$ | * | 0.21 | $1.14 \times 10^{-5}$ | * |
| PPA to PCU | -0.25 | $5.70 \times 10^{-8}$ | * | -0.23 | $2.31 \times 10^{-6}$ | * |
| HC to PPA | -0.24 | $2.06 \times 10^{-7}$ | * | -0.21 | $1.14 \times 10^{-5}$ | * |
| HC to HC | -0.03 | 0.562 | | -0.04 | 0.383 | |
| HC to PCU | 0.01 | 0.855 | | -0.03 | 0.560 | |
| PCU to PPA | 0.26 | $1.08 \times 10^{-8}$ | * | 0.21 | $1.14 \times 10^{-5}$ | * |
| PCU to HC | 0.09 | 0.053 | | 0.03 | 0.532 | |
| PCU to PCU | 0.08 | 0.104 | | 0.06 | 0.237 | |
| **Modulation** | | | | | | |
| PPA | -0.27 | $6.14 \times 10^{-9}$ | * | -0.24 | $3.34 \times 10^{-7}$ | * |
| HC | -0.09 | 0.069 | | -0.08 | 0.098 | |
| PCU | -0.15 | 0.001 | . | -0.14 | 0.005 | . |
| **Input** | | | | | | |
| PPA | 0.32 | $2.66 \times 10^{-12}$ | * | 0.31 | $3.66 \times 10^{-11}$ | * |

The table presents the correlation coefficients ($\rho$) and p-values for each connection's relationship with memory performance and PACC5 test. To indicate the amplitude of the effect, the significance column uses the following symbols: '*' for $p < 0.05$ with $|\rho| > 0.2$ and '.' for $p < 0.05$ but $|\rho| \leq 0.2$. P-values were adjusted for false discovery rate (FDR) using the Benjamin-Hochberg procedure. Self-connection of parahippocampal place area (PPA) was excluded from this analysis as it was not considered to be correlated with memory performance. Abbreviations: PPA (parahippocampal place area), HC (hippocampus), and PCU (precuneus)

**Supplementary table 4 The results from generalised additive model of hippocampal volume onto the connectivity of interest**

| Connectivity | Hippocampal volume |
|---|---|
| **Intrinsic** | |
| PPA to HC | 0.446 |
| PPA to PCU | 0.649 |
| HC to PPA | 0.446 |
| PCU to PPA | 0.649 |
| **Modulatory** | |
| PPA | 0.446 |
| **Input** | |
| PPA | 0.749 |

The table displays the p-values (FDR-corrected using the Benjamin-Hochberg procedure) for each term in the models. The contribution of hippocampal volume is not significant on any connectivity. Abbreviation: PPA, parahippocampal place area; HC, hippocampus; PCU, precuneus.

**Supplementary Figure 1 Relationship between the effective connectivity and memory performance.**

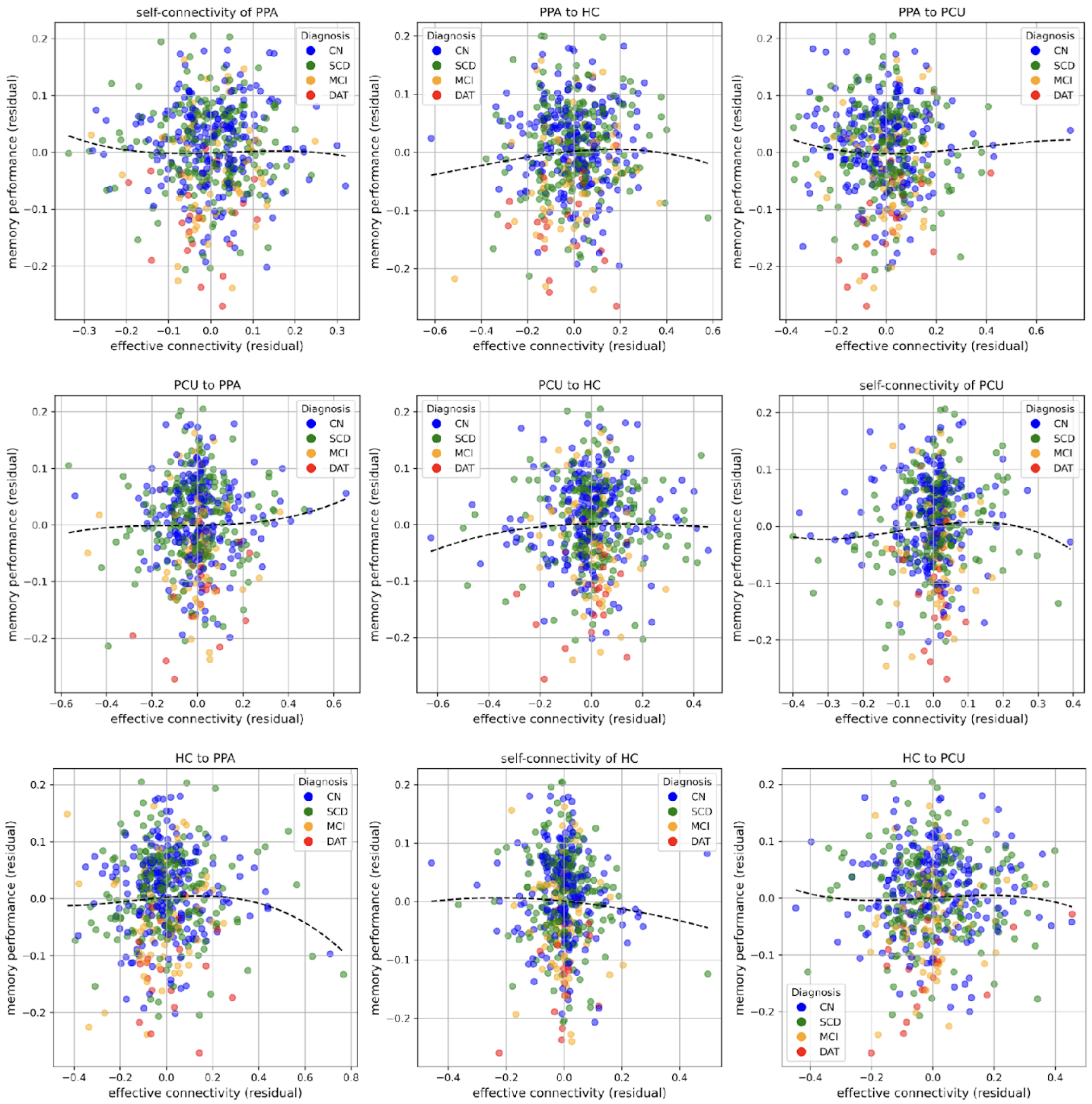


The plots illustrate the spline relationships between effective connectivity and memory performance. The terms are presented as residuals, adjusted for age, sex, education, and other connectivity estimates. Data points are color-coded or highlighted according to diagnoses: CN (cognitively normal), SCD (subjective cognitive decline), MCI (mild cognitive impairment), and DAT (dementia of Alzheimer's type). A' denotes memory performance. Abbreviations: PPA (parahippocampal place area), HC (hippocampus), and PCU (precuneus)
